# Neutron capture enhances dose and reduces cancer cell viability in and out of beam during helium and carbon ion therapy

**DOI:** 10.1101/2023.12.03.569810

**Authors:** Nicholas Howell, Ryan J. Middleton, Frederic Sierro, Naomi A. Wyatt, Andrew Chacon, Benjamin H. Fraser, Keith Bambery, Elle Livio, Christopher Dobie, Joseph J. Bevitt, Justin Davies, Anthony Dosseto, Daniel R. Franklin, Ulf Garbe, Susanna Guatelli, Ryoichi Hirayama, Naruhiro Matsufuji, Akram Mohammadi, Karl Mutimer, Louis M. Rendina, Anatoly B. Rosenfeld, Mitra Safavi-Naeini

**Affiliations:** Australian Nuclear Science and Technology Organisation, Lucas Heights, NSW, Australia; Wollongong Isotope Geochronology Laboratory, School of Earth, Atmospheric and Life Sciences, University of Wollongong, Wollongong, NSW, Australia; School of Electrical and Data Engineering, University of Technology Sydney, Ultimo, NSW, Australia; Centre for Medical Radiation Physics, University of Wollongong, Wollongong, NSW, Australia; National Institutes for Quantum Sciences and Technology, Chiba, Japan; School of Chemistry, The University of Sydney, Sydney, NSW, Australia; The University of Sydney Nano Institute, Sydney, NSW, Australia

## Abstract

**Purpose:** Neutron Capture Enhanced Particle Therapy (NCEPT) is a proposed augmentation of charged particle therapy which exploits thermal neutrons generated internally, within the treatment volume via nuclear fragmentation, to deliver a biochemically targeted radiation dose to cancer cells. This work is the first experimental demonstration of NCEPT, performed using both carbon and helium ion beams with two different targeted neutron capture agents (NCAs).

**Materials and Methods:** Human glioblastoma cells (T98G) were irradiated by carbon and helium ion beams in the presence of NCAs, [^10^B]-BPA and [^157^Gd]-DOTA-TPP. Cells were positioned within a PMMA phantom either laterally adjacent to, or within, a 100×100×60 mm spread out Bragg peak (SOBP). The impact of NCAs and location relative to the SOBP on the cells was measured by cell growth and survival assays in six independent experiments. Neutron fluence within the phantom was characterised by quantifying the neutron activation of gold foil.

**Results:** Cells placed inside the treatment volume reached 10% survival by 2 Gy of C or 2-3 Gy of He in the presence of NCAs compared to 5 Gy of C and 7 Gy of He with no NCA. Cells placed adjacent to the treatment volume showed a dose-dependent decrease in cell growth when treated with NCAs, reaching 10% survival by 6 Gy of C or He (to the treatment volume), compared to a no detectable effect on cells without NCA. The mean thermal neutron fluence at the centre of the SOBP was approximately 2.2×10^9^ n/cm2/Gy(RBE) for the carbon beam and 5.8×10^9^ n/cm2/Gy(RBE) for the helium beam and gradually decreased in all directions.

**Conclusions:** The addition of NCAs to cancer cells during C and He beam irradiation has a measurable impact on cell survival and growth *in-vitro*. Through the capture of internally generated neutrons, NCEPT introduces the concept of a biochemically targeted radiation dose to charged particle therapy. NCEPT enables the established pharmaceuticals and concepts of neutron capture therapy to be applied to a wider range of deeply situated and diffuse tumours, by targeting this dose to micro-infiltrates and cells outside of defined treatment regions. These results also demonstrate the potential for NCEPT to provide an increased dose to tumour tissue within the treatment volume, with a reduction in radiation doses to off target tissue.

## Introduction

Radiation therapy aims to treat cancer by delivering a therapeutic dose to the entire tumour while minimising radiation exposure to healthy tissue. Of the various radiation therapy methods, charged particle therapy (either proton or heavy ion therapy^1–4)^ is amongst the most effective at achieving this goal. This is because charged particles can deliver a highly conformal radiation dose with a small number of treatment fractions, even to deep tumours, due to the physical characteristics of an ion’s Bragg peak. The energy loss of a charged particle is inversely proportional to the square of its velocity, resulting in the majority of its kinetic energy being deposited at the end of its track just before it comes to rest.

The effectiveness of charged particle therapy is limited by the accuracy with which the tumour can be delineated during pre-irradiation PET/CT or PET/MR imaging, since unresolvable micro-infiltrates and micro-metastasis cannot be deliberately targeted for irradiation. Furthremore, while the entrance dose - the dose received by tissue through which the beam passes before reaching the tumour - is lower than for photon therapy, it remains substantial and must be carefully limited during treatment planning. Additionally, inelastic collisions between accelerated charged particles and the target result in a range of nuclear fragments, including lighter ions and neutrons which deposit additional dose in surrounding tissues^2^. The neutron component of this radiation field extends almost isotropically around the target volume, and due to scattering within the patient, the neutrons rapidly lose kinetic energy and approach thermal equilibrium with their surroundings^4,5^.

An extension of charged particle therapy that utilises the neutron component of the radiation field within the patient via neutron capture was proposed in 2018 (Figure 1). Neutron capture therapy (NCT) has been previously used for cancer treatment by systemic administration of a neutron capture agent (NCA) to the patient and irradiating the target tissue with an external neutron source. In NCT, thermal neutrons are captured by isotopes with high thermal neutron capture cross-sections, releasing high-LET charged particles that damage cancer cells^6–9^. However, the use of an external neutron source limits the depth at which sufficient neutrons can be delivered to the target without causing excessive radiation induced proximal tissue injury.

**Figure 1:**
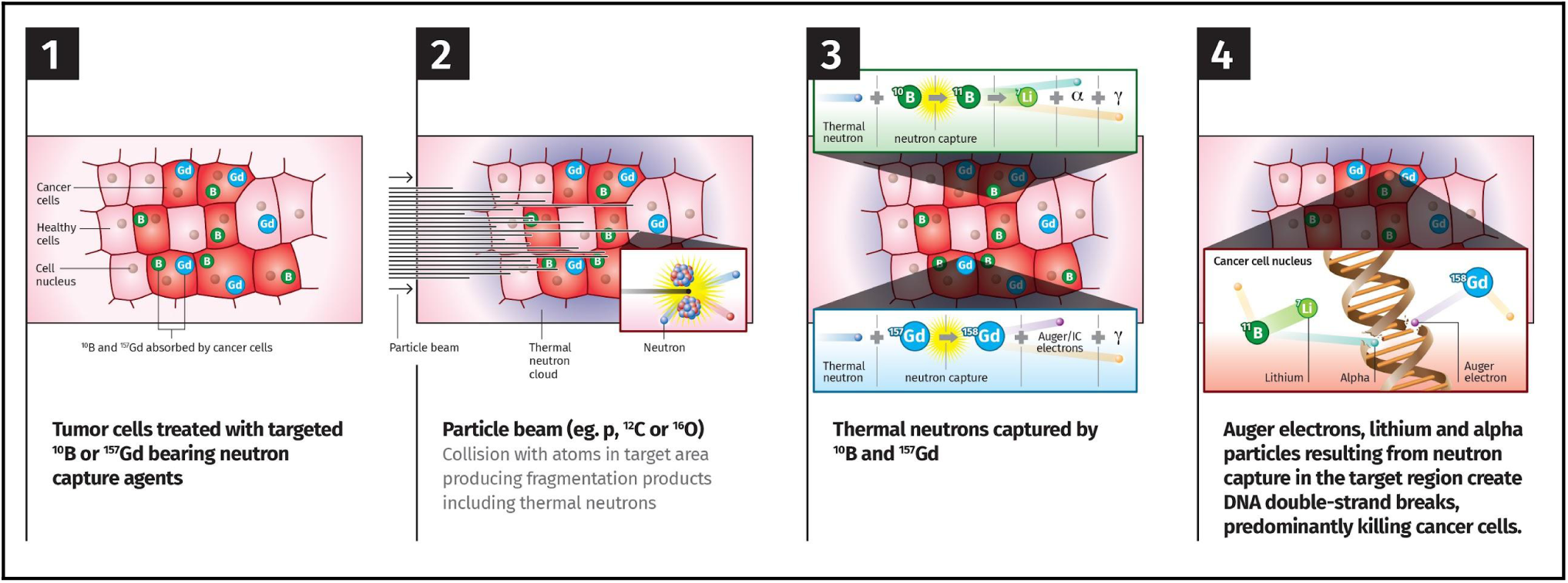
Physics of NCEPT: As ions in the beam traverse tissues proximal to the target, some undergo nuclear interactions with target matter, creating a variety of nuclear fragmentation products, including neutrons, which disperse in the body and thermalise. Target-specific agents deliver ^10^B or ^157^Gd payloads to cancer cells, where thermal neutrons are captured resulting in emission of short-range (4-8 μm for ^10^B, 10-20 nm for ^157^Gd), high-LET particles. While the particle beam treats the bulk of the tumour, the neutrons released are captured anywhere that NCAs are present - inside and outside the primary treatment volume - that would have otherwise remained unirradiated.

Neutron capture enhanced particle therapy (NCEPT) aims to combine the spatial precision of charged particle therapy with the precise biochemical targeting of NCT. NCEPT offers two major benefits compared to conventional charged particle therapy:

1. The dose to the target volume can be increased relative to the surrounding tissue (including the entrance region) due to the additional contribution of the neutron capture dose; and
2. Since the neutron field is generated internally and extends beyond the primary treatment volume, a biochemically targeted NCA which preferentially concentrates in cancer cells can deliver a therapeutically useful dose even to small satellite lesions beyond the primary target volume.

The theoretical feasibility of this method has previously been established via Monte Carlo simulations, demonstrating that a representative proton or heavy ion therapy treatment plan will generate a sufficient thermal neutron field to deliver an additional dose of at least 10% with achievable tissue concentrations of both ^10^B and ^157^Gd-based NCAs^10^. While dose enhancement has previously been reported experimentally during proton therapy in the presence of natural boron by Cirrone et al. and Bláha et al. and attributed to proton-^11^B fusion^11,12^, it now appears that the vast majority of this effect is actually due to neutron capture by ^10^B^13,14^, which has a natural isotopic abundance of 19.8%^15^, rather than proton-^11^B fusion (with Jacobsen et al. reporting that the number of high-LET particles produced by neutron capture outnumber those produced via fusion by a ratio of ∼4000±700:1^13^). This hypothesis is especially supported strongly by the failure of the result to be reproduced experimentally in the absence of a surrounding phantom (which is necessary to generate a neutron field, but should not be necessary for proton-^11^B fusion)^16,17^. Therefore, if neutron capture is the dominant cause of dose enhancement, then the effect should also be observed during helium or carbon ion therapy in the presence of enriched ^10^B (or other high neutron capture cross-section isotopes such as ^157^Gd), where proton fusion plays no role whatsoever.

In this paper, the NCEPT principle is established *in vitro* using two NCAs irradiated by helium and carbon ions. The first, 4-borono-L-phenylalanine ([^10^B]BPA, Figure 2(a)) is an unnatural amino acid which has been used clinically in conventional neutron capture therapy for more than two decades^18^. BPA is transported primarily through amino acid transporter LAT1, and, to a lesser extent, LAT2 and ATB(0+)^19^. LAT1 is a plasma membrane amino acid transporter that is expressed in a variety of cancer tissues and high expression is associated with poor prognosis^20^. ^10^B has a high capture cross-section for thermal neutrons (∼3,800 barns). Upon capturing a thermal neutron, an excited ^11^B* nucleus is formed which immediately fissions to release two high LET ions - an alpha particle and a + ^7^Li nucleus^21^, with ranges of approximately 4.1 and 7.7 μm in tissue, respectively.

**Figure 2:**
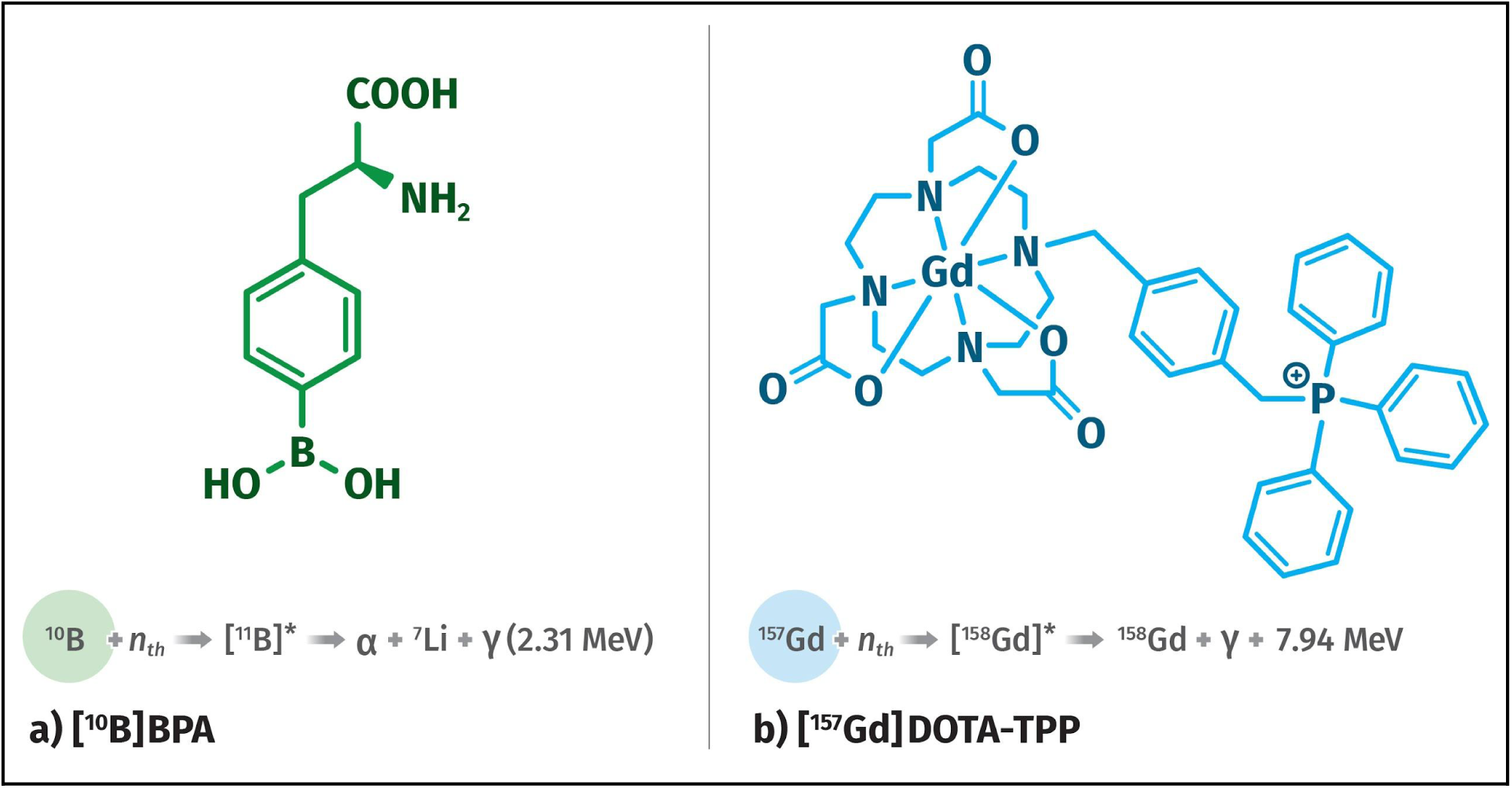
Neutron capture agents (NCAs) used in this work and their respective thermal neutron capture reactions; (a) ^10^B-enriched 4-borono-L-phenylalanine; (b) ^157^Gd-enriched 2,2′,2′′-[10-{2-[(triphenylphosphonio)methyl]benzyl}-1,4,7,10-tetraazacyclododecane-1,4,7-triyl ]triacetatogadolinium(III) trifluoroacetate^31,32^

The second NCA is the novel compound 2,2′,2′′-[10-{2-[(triphenylphosphonio)methyl]benzyl}- 1,4,7,10-tetraazacyclododecane-1,4,7-triyl]triacetatogadolinium(III) trifluoroacetate ([^157^Gd]DOTA-TPP, Figure 2(b)), which uses the mitochondria-targeting moiety triphenylphosphonium (TPP) to accumulate ^157^Gd in the inner mitochondrial membrane (IMM). TPP is particularly efficient at accumulating in negatively charged membrane compartments^22^. This compound preferentially accumulates in cancer cells due to the increased mitochondrial function in many cancers^23–25^. ^157^Gd is a stable isotope of gadolinium with an extremely high thermal neutron capture cross-section (∼2.55 x 10^5^ barns). Neutron capture by ^157^Gd results in the formation of an excited ^158^Gd* nucleus; its subsequent de-excitation results in the emission of an average of 5 Auger and 0.69 internal conversion electrons, plus an average of 1.8 high energy prompt gamma photons and one recoil ^158^Gd nucleus^26–29^. The high LET Auger electrons have a range of the order of 10 to 20 nm; accumulation of damage to the IMM will, in most cases, lead to apoptosis^30^.

Demonstrated below is the effect of NCEPT on human glioblastoma multiforme cell culture when irradiated with helium and carbon ion beams, and emphasises the potential significance of including neutron capture to the treatment volume itself as well as the surrounding area in the overall treatment plan.

### Methods and Materials

The experiments described in the following sections have three primary objectives:

1. Confirming the magnitude and distribution of the predicted thermal neutron fluence in and around the primary radiation field (target volume);
2. Determining whether the results predicted via the previous simulation study would translate to effective attenuation of cancer cell proliferation in vitro, using ^4^He and ^12^C ion beams and two neutron capture agents; and
3. Determining the effect of neutron capture on cancer cells outside of the primary radiation field but still within the thermal neutron field generated by the heavy ion irradiation.

The experimental configuration is shown in Figure 3. All experimental measurements were performed using the Heavy Ion Medical Accelerator in Chiba (HIMAC) biological beamline at the National Institute for Quantum Science and Technology (QST) in Japan.

**Figure 3:**
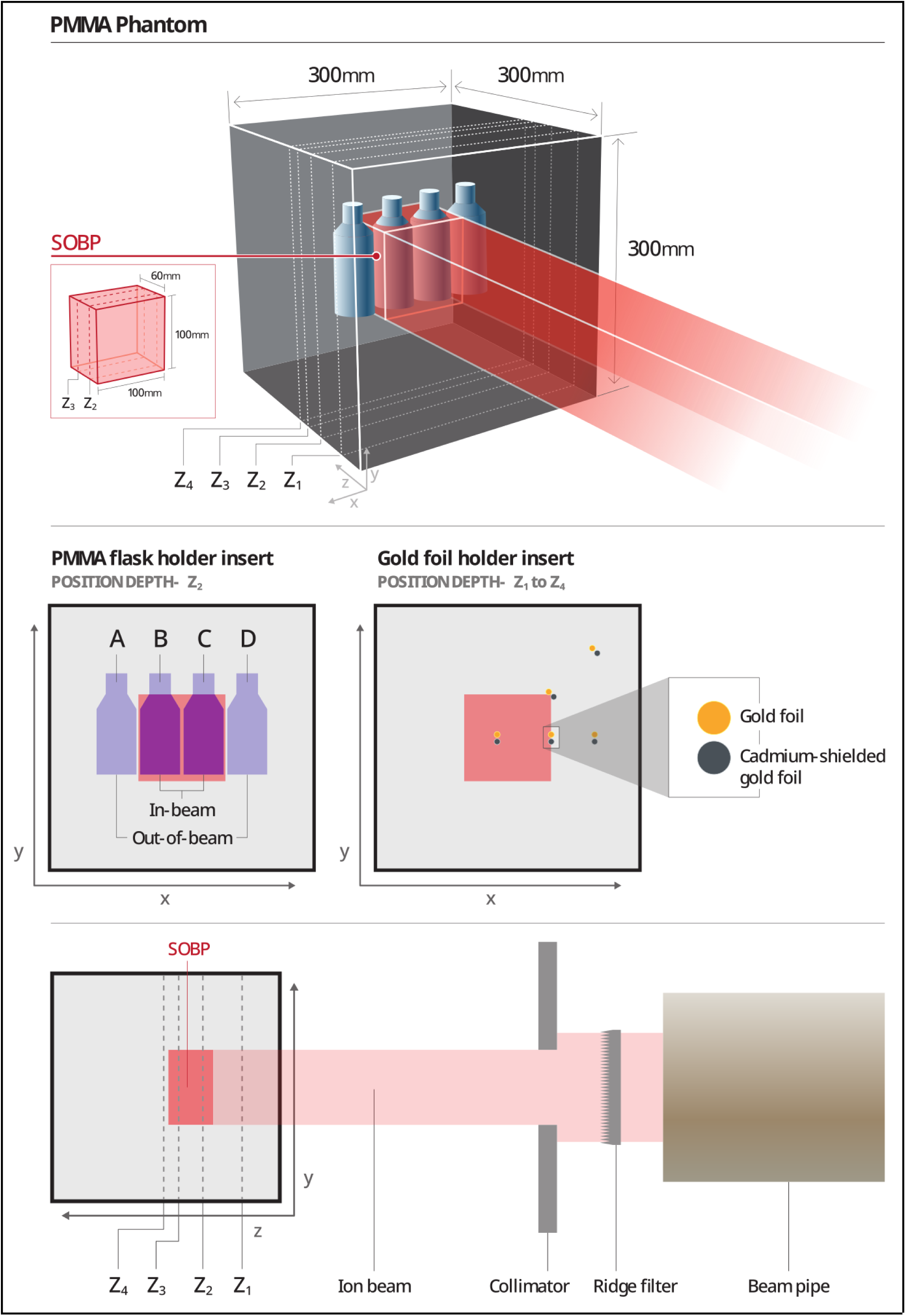
Experimental configuration; irradiation positions for flask irradiation and neutron fluence measurements; positions **A** and **D** are out of beam; **B** and **C** in-beam. The spread out Bragg peak (SOBP) extends 60 mm from a depth of 80 mm to 140 mm (in the z dimension), with a 100 mm × 100 mm cross-sectional area. Gold activation measurements of thermal neutron fluence were made at depths of Z_1_ = 48 mm (proximal to the SOBP), Z_2_ = 100 mm (inside the SOBP), Z_3_ = 133 mm (inside the SOBP) and Z_4_ = 153 mm (distal to the SOBP). For further detail see Supplementary Material Figure 1.

### Phantom and irradiation conditions

A multi-layer cubic PMMA phantom (ρ=1.19 g/cm^3^) was assembled with total dimensions of 300×300×300 mm^3^. PMMA slab inserts were constructed with either paired circular indentations for holding gold foils, or with single or double cut-outs for holding T25 flasks filled with medium (Figure 3). The cut-outs for the T25 flasks are positioned so that the cell layer (at the bottom of the flasks) was located at depth Z_2_ (100 mm) in PMMA, perpendicular to the beam.

For both carbon and helium ion beams, a 100×100×60 mm^3^ SOBP was used, corresponding to a depth range of 80-140 mm in PMMA. The SOBPs were generated via passive scattering of monoenergetic 150 MeV/u and 290 MeV/u helium and carbon ion beams, respectively, with the distal edges (i.e. the maximum depths) of the SOBPs corresponding to the energies of the respective monoenergetic ion beams.

### Thermal neutron fluence measurements

The differential neutron activation method for thermal neutron fluence measurement was validated beforehand via Monte Carlo simulations (as described in the Supplementary Material Section 1; physics models are detailed in Supplementary Material Table 1 and the results obtained via simulation are compared with ground truth in Supplementary Material Tables 3 and 4).

The neutron detector consists of pairs of thin disc foils of pure ^197^Au (average mass=12.075 mg, *ρ*=19.32 g.cm^-3^, nominal *r*=3 mm, nominal *T*=0.02218 mm); one disc is bare while the other is covered by a 1 mm layer of cadmium on both sides to absorb thermal neutrons. Paired foils were placed at different positions in a 5 mm thick PMMA layer, which was then inserted into the phantom at depths of 48 mm (proximal to the SOBP), 100 mm and 133 mm (both inside the SOBP), and 153 mm (distal to the SOBP). At each depth, foil pairs were placed at lateral offsets of -12 mm (rather than zero, so as to avoid overlap with successive depths), +50 mm and +100 mm from the central axis of the phantom. Additional foil pairs were placed on a radius at a 45° offset in the transaxial plane, with the pairs centred at points 70.7 mm and 154 mm from the centre of the phantom. Therefore, each plate included a total of 5 pairs of foils. A schematic diagram of the plate outline is shown in Figure 3 (specific locations are described in Supplementary Material Table 2). Successive plates are stacked and progressively rotated by 90° to avoid attenuation of the primary particle beam by preceding foils.

Total doses of 70 Gy(RBE) and 35 Gy(RBE) were used for irradiation with helium and carbon, respectively. The resulting gold foil activities were measured over a period of 1 h using a high purity germanium (HPGe) detector (Canberra GC-2020), and the number of photons detected within the 411 keV gamma peak was converted to thermal neutron fluence using (1):

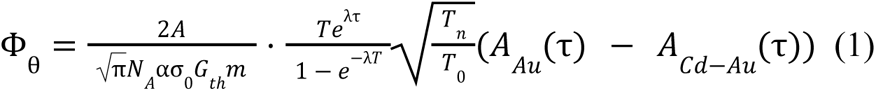

where *τ* is the time after irradiation that the measurements are performed (zero if the activity is decay-corrected), *A_Au_ (τ)* is the post-irradiation activity of the bare gold foil (due to all neutron captures) at *t = τ*, *A_Cd-Au_*(*τ*) is the activity of the cadmium-shielded foil at *t = τ* , *A* is the mass number of gold (assumed to be 196.9666), *N_A_* is Avogadro’s Number, *α* is the abundance of the target isotope ^197^Au (assumed to be 1), *σ_0_* is the thermal neutron cross-section of ^197^Au (9.87×10^-23^ cm^2^), *G_th_* is the self-shielding ratio, *m* is the mass of the sample, *λ* is the decay constant of ^198^Au, *T* is the duration of irradiation, *T_n_*is the sample temperature, and *T_0_* is the reference temperature (both are 293 K in this case)^33,34^. The resulting fluence estimate is divided by the biological dose to provide the flux per unit dose (n/cm^2^/Gy(RBE))^34,35^. A full derivation of the self-shielding ratio for the experimental configuration is provided in the Supplementary Material Section 2.

### Cell Cultures

The human glioblastoma multiforme cell line, T98G, was sourced from the Japan Cancer Research Resources Bank. T98G was maintained as a sub-confluent culture in a humidified incubator at 37℃ with 5% CO_2_ grown in Eagles Modified Essential Media (EMEM) (Gibco) supplemented with 10% foetal bovine serum plus penicillin and streptomycin antibiotics.

### Neutron Capture Agents (NCAs)

Two neutron capture agents were used in this study - [^10^B]BPA and [^157^Gd]DOTA-TPP. The chemical structure of these compounds and the relevant neutron capture reactions are shown in Figure 2. [^10^B]BPA (>98.4% ^10^B) was purchased from Interpharma Praha, a.s., (Prague, Czech Republic). [^157^Gd]DOTA-TPP (>88% ^157^Gd) was synthesised using a two-step procedure adapted from Morrison *et al*^32^. In brief, commercially available DO3A-*tert*-butyl ester (Macrocyclics) was alkylated with 4-(bromomethyl)benzyl)triphenyl phosphonium bromide under basic conditions to yield a solid which was then reacted, without further purification, with trifluoroacetic acid (TFA) which cleaved the *tert*-butyl ester protecting groups. The excess TFA was then removed by evaporation under reduced pressure, and the resulting crude material was purified by reverse-phase preparative HPLC to give the DOTA-TPP ligand as white solid. The purified DOTA-TPP ligand was then dissolved up in water and treated with Gd-157 enriched (88%) gadolinium oxide. The final product was obtained, after filtration and lyophilisation, as white solid and gives HPLC (QC), elemental analysis and mass spectrometry data consistent with that expected for the gadolinium-157 enriched metal complex ([^157^Gd]DOTA-TPP) (see Supplementary Material Section 3 for full details).

### Cell irradiation conditions

Irradiations of T98G cultures were conducted in 80-90% confluent T25 tissue culture flasks. Approximately 24 h prior to irradiation, the cultures were treated with either 500 μM [^10^B]BPA, 500 μM [^157^Gd]DOTA-TPP or a vehicle control.

Carbon and helium ion beams were configured to deliver nominal dose rates of ∼1 Gy/min, with exact exposure times adjusted based on nightly dose rate measurements at the beginning of each experimental session. Total doses of 0, 2, 3, 4, 6 and 8 Gy were used for irradiation of the NCA-treated and control flasks (control flasks received an additional dose of 10 Gy) for both ions. Immediately prior to irradiation, the flasks were topped up to the base of the neck with either PBS or EMEM. Flasks were irradiated at positions B and C, with the cell layer at a depth of 100 mm within the PMMA phantom (Figure 3), at room temperature, either in pairs or alongside a sham flask containing only water. Following irradiation (5-10 min from beam-off) the media was vacuum aspirated, the T98G monolayer detached and processed as described in the following sections for both cell growth (resazurin) and/or clonogenic assays.

To evaluate the efficacy of neutron capture events occurring outside of the primary beam field, flasks were prepared as above, with either 500 µM [^10^B]BPA, [^157^Gd]DOTA-TPP or diluent control. The flasks were placed at positions A and D within the phantom while B and C received a sham water flask. Total doses of 0, 1, 3, 6 and 10 Gy of carbon and helium ions were delivered by the primary beam to the two sham flasks.

The efficacy of [^10^B]BPA and [^157^Gd]DOTA-TPP as a function of concentration was investigated by fixing the physical dose at 3 Gy and irradiating T98G cultures under progressively decreasing concentrations of NCA. 3 Gy was chosen as it is approximately equal to the LD30 for T98G cells in response to direct irradiation with the C or He particle beam. A dilution series was created for each compound in EMEM (500, 250, 100, 50, 25, 10, 5, 2.5 and 1 μM) plus a vehicle control. Cells were incubated with NCA for approximately 24 h and irradiated, as above, with 3 Gy of carbon ions or helium ions at a dose rate of ∼1 Gy/min and immediately processed as described below. A 0 Gy (minimum response) and 10 Gy (maximum response) control were also included for each NCA treatment group and vehicle control.

### Cell growth assay

The resazurin reduction based cell proliferation assay was the preferred method chosen for measuring the impact of NCEPT *in vitro*. It has several advantages over clonogenic assay under these conditions (especially extremely limited beamtime), including increased throughput, straightforward data interpretation and observability of growth kinetics including lag and plateau phases. The use of a metabolic reduction as the readout means that the assay takes into account all viable cells within a population, including those who have left the cell cycle due to radiation impact (e.g. senescence, quiescence). This distinguishes this approach from the traditional clonogenic assay and leads to the production of a more conservative, but overall representative, view of the cellular impact in a non-uniform mixed radiation field. This method was employed for quantifying in-beam and out-of-beam dose response as well as the (in-beam) NCA concentration response with a fixed ion dose.

Following irradiation, the cell monolayer was detached with trypsin and the resulting cell suspensions were adjusted to 2,500 cells/mL. 200 μL of this suspension was pipetted into the designated wells of 7 × 96 well, black wall, clear bottom plates to give 500 cells/well. Plates were incubated at 37 ℃ with 5% CO_2_. One plate was assayed every 24 h for viable cell mass using resazurin assay as follows^36^. A 1% w/v stock solution of resazurin sodium salt was prepared in water. The stock was diluted 250 times in phosphate-buffered saline (PBS) and warmed to 37℃ in a water bath. All media was removed from the plate, replaced with resazurin-PBS and incubated for 60 min at 37℃. The resulting conversion of resazurin to resorufin was measured by recording the fluorescence at excitation wavelength 555 nm and emission wavelength 585 nm at 37℃. The resulting relative fluorescence units (RFU) were plotted against time (days) to quantify the proliferation rates and growth of T98G cultures following irradiation with and without a NCA^37^ (Supplementary Material Figure 2 and Figure 3). A time point was selected from this data in which all treatments were in as close to an exponential growth phase as possible (*i.e.* 144 h) and the RFU re-plotted against log dose (Gy), with 95% confidence intervals, for each well of cells treated with either no NCA, [^10^B]BPA or [^157^Gd]DOTA-TPP and irradiated with both carbon and helium ion beams. Lethal dose, 50% (LD50) was determined by fitting a sigmoidal log-inhibitor vs. response model to this data and percent response (F) was calculated from this LD50 and the Hill-slope (H) using the relationship:

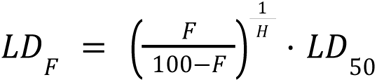

### Clonogenic Assay

Clonogenic assay was also employed as a secondary method for evaluation of in-beam cell survival in response to ion irradiation with and without the presence of NCAs.

Following irradiation, flasks were trypsinised and T98G cells were seeded into Petri dishes (Falcon 100 mm Cell Culture Dish, Corning) with concentrations listed in Supplementary Material Table 7. The cells were then incubated at 37°C with 5% CO_2_ for 12 days (approximately 10 doubling times^38^). The resulting cell cultures were washed with 5 mL of PBS, fixed in 5 mL of 10% neutral buffered formalin, washed with distilled water and stained with crystal violet to highlight the nucleus and aid with colony identification. Each dish was then left to dry and digitised using the ImageQuant LAS 4000 (GE) imaging system^39^. The number of colonies (>50 cells) in each dish was estimated using the OpenCFU software^40^ (See Supplementary Material Figure 4). Survival fraction was then calculated by dividing the number of colonies by the initial seeding density.

The mean survival fraction was plotted on a logarithmic scale against dose, with 95% confidence intervals, for each population of cells treated with either no NCA, [^10^B]BPA or [^157^Gd]DOTA-TPP and irradiated with both carbon and helium ion beams.

The survival fraction of cells irradiated by photons or charged particles generally follows a linear-quadratic function of dose, according to the relation

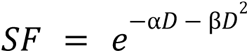

The ɑ and β parameters were computed using a least-squares error minimisation algorithm for each case (Supplementary Material Table 8), and the 1% and 10% survival fractions estimated for each case.

## Results

### Thermal neutron fluence measurements

The thermal neutron fluences at each point of measurement resulting from irradiation of the PMMA target with helium ions are plotted in space relative to the SOBP in Figure 4(a) and listed in Supplementary Material Table 5; corresponding results for carbon are shown in Figure 4(b) and Supplementary Material Table 6. The mean thermal neutron fluence at the centre of the SOBP is approximately 2.2×10^9^ (±1.4×10^8^) n/cm^2^/Gy(RBE) for the carbon beam and 5.8×10^9^ (±5.3×10^7^) n/cm^2^/Gy(RBE) for the helium beam.

**Figure 4:**
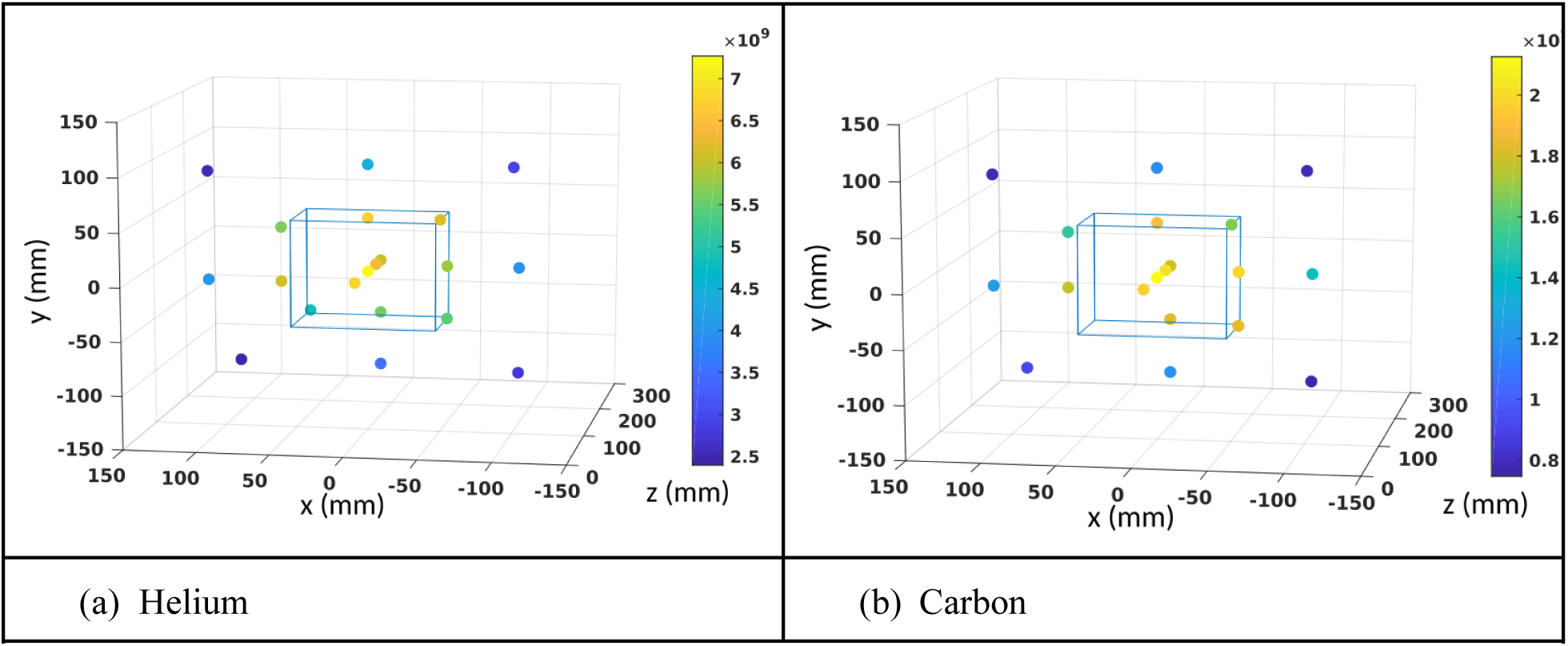
Neutron fluence per unit dose (n/cm^2^/Gy(RBE)) at the evaluated locations for both beam types. The wireframe rectangular prism represents the SOBP region.

### In vitro evaluation of in-beam NCEPT

The fractions of T98G cells which remain viable following in-beam irradiation with helium and carbon ions are plotted as a function of increasing dose in Figure 5. Three treatment conditions were used for each ion: either no NCA, 500 μM [^10^B]BPA or 500 uM [^157^Gd]DOTA-TPP.

**Figure 5:**
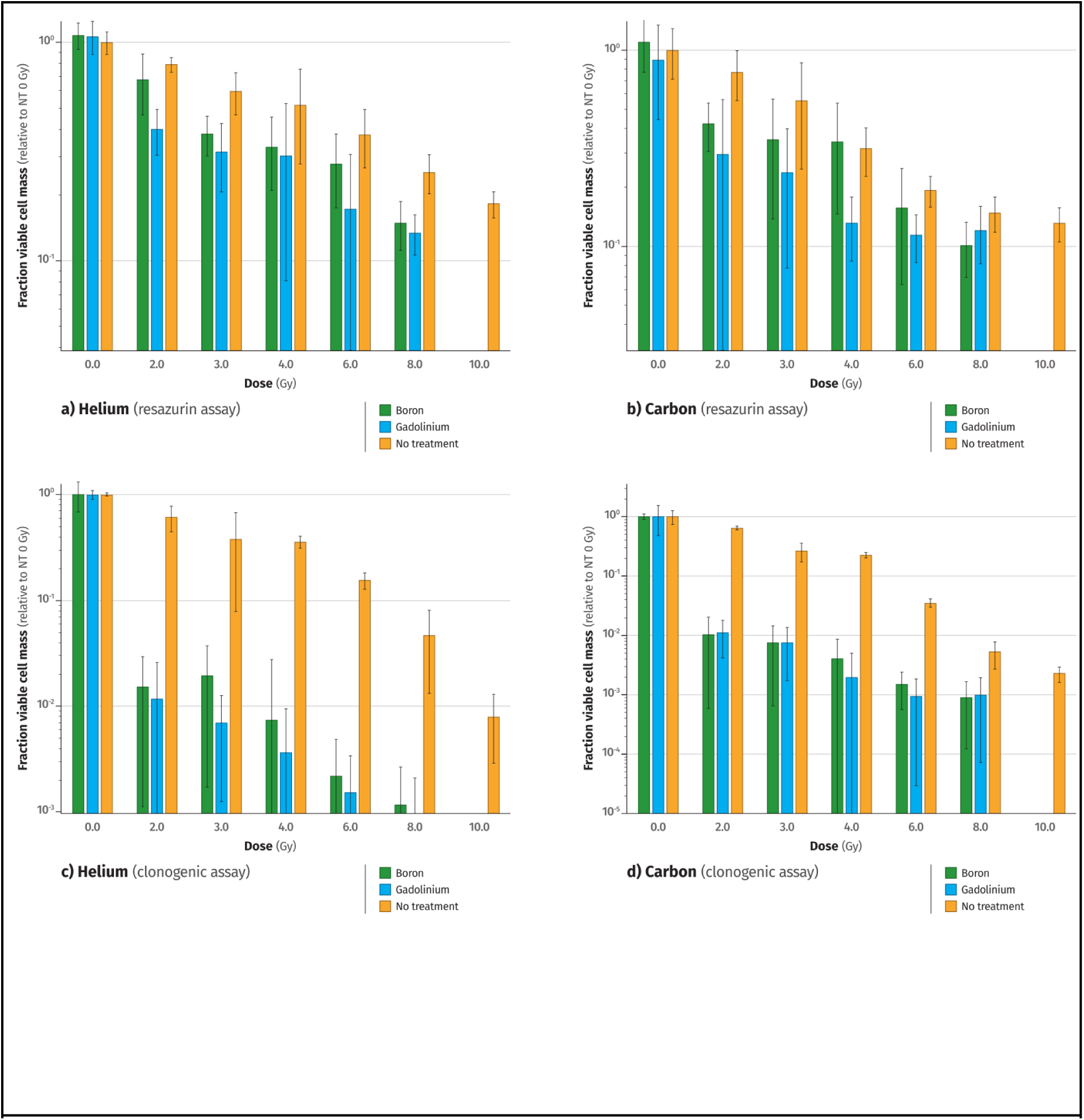
Dose-dependent effect of neutron capture agents (NCAs) on cell survival. Cell survival fraction (normalised to no-treatment, 0 Gy) for T98G cells positioned in the centre of the spread out Bragg peak region and irradiated with helium ions (a & c) or carbon ions (b & d) across a range of physical radiation doses, following incubation with 500 μM of either neutron capture agent (NCA), in triplicate. Cell survival was evaluated using resazurin (a & b) and clonogenic (c & d) assays. Bars represent the mean fraction viable cell mass of three independent experiments for the clonogenic, while the resazurin assay shows representative data of six independent experiments. T-bars represent ±2 standard deviations.

**Figure 6:**
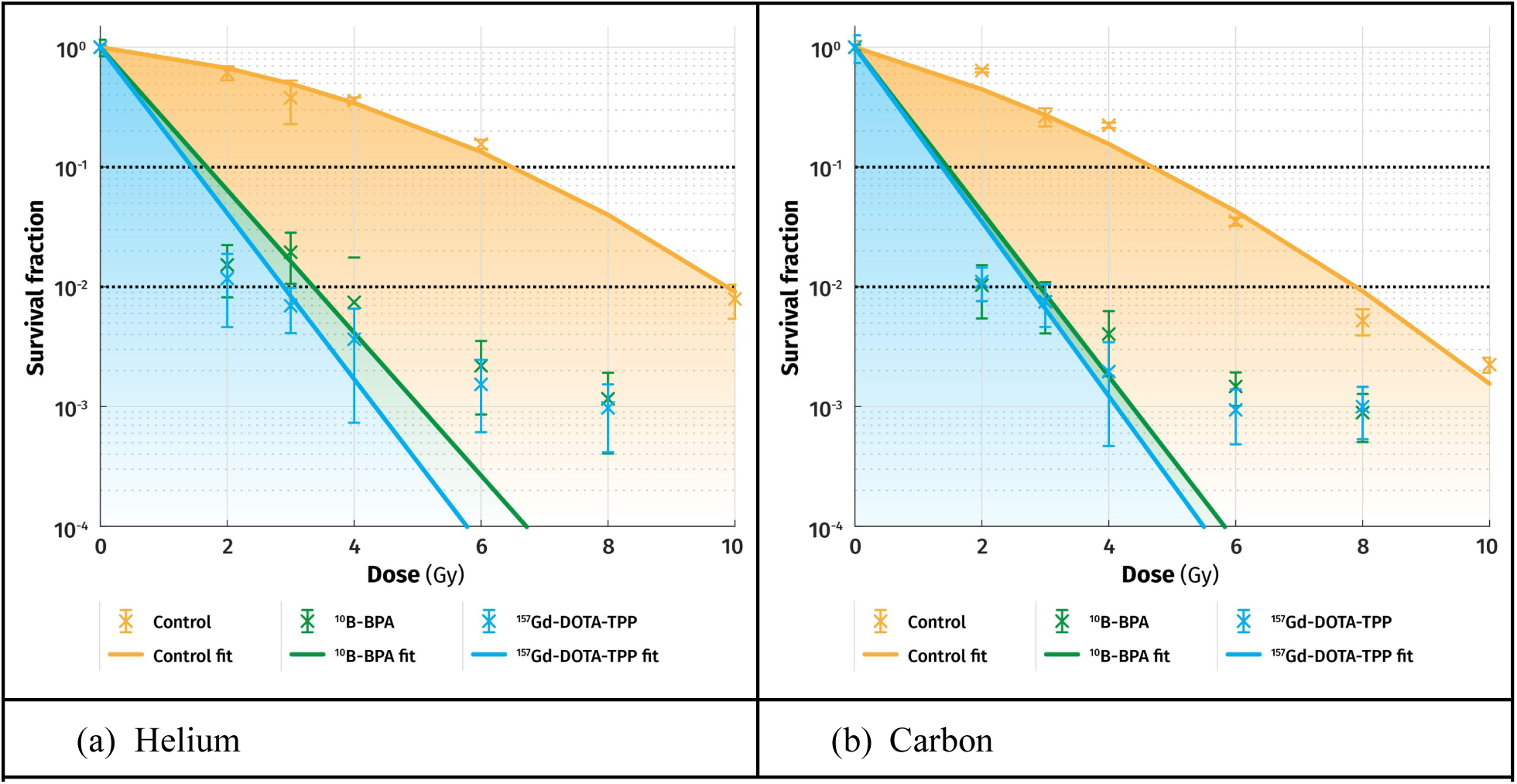
Mean survival fractions as a function of dose (95% confidence intervals) of three independent experiments, for each population of cells treated with no NCA, [^10^B]BPA or [^157^Gd]DOTA-TPP and irradiated with (a) helium and (b) carbon ion beams. The ɑ and β parameters of the linear quadratic model were computed using a least-squares fitting process algorithm for each case (Supplementary Material Table 8); 10% and 1% survival thresholds are shown as horizontal dashed lines.

The corresponding LD50 and LD90 doses estimated from the resazurin assay results, and 10% and 1% survival fraction doses obtained from the clonogenic assay results are shown in Table 1.

**Table 1.**
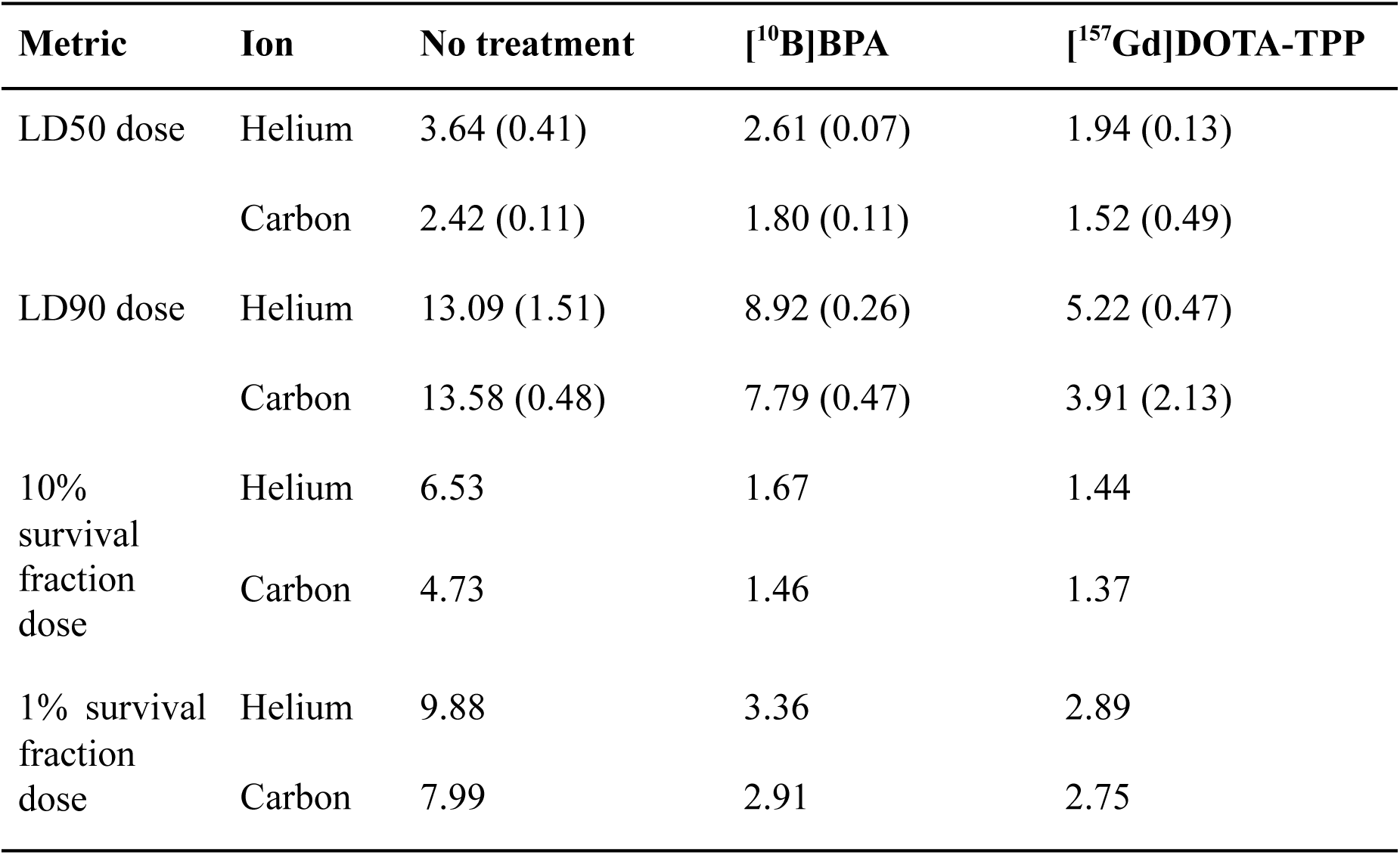
Mean LD50 and LD90 of six independent experiments (resazurin cell growth assay; see Figures 5(a) and 5(b)), and 10% and 1% survival fractions of three independent experiments (clonogenic assay, with survival thresholds estimated based on the fitted linear quadratic model curves in Figure 6). T98G cells were placed at a depth of 110 mm in a 300 mm cubic PMMA phantom and irradiated in-beam, at the centre of the SOBP60 (Figure 3) with helium or carbon ions, either with NCA present (control), 500 uM [^10^B]BPA or 500 uM [^157^Gd]DOTA-TPP.

### In vitro evaluation of out-of-beam NCEPT

The fractions of T98G cells which remain viable following out-of-beam irradiation with helium and carbon ions are plotted as a function of increasing dose to the primary target in Figure 7. Cells were positioned within the phantom, immediately adjacent to the primary target volume and not in the path of the primary beam (at positions A and D shown in Figure 2). Again, three treatment conditions were used for each ion: either no NCA, 500 uM [^10^B]BPA or 500 uM [^157^Gd]DOTA-TPP.

**Figure 7:**
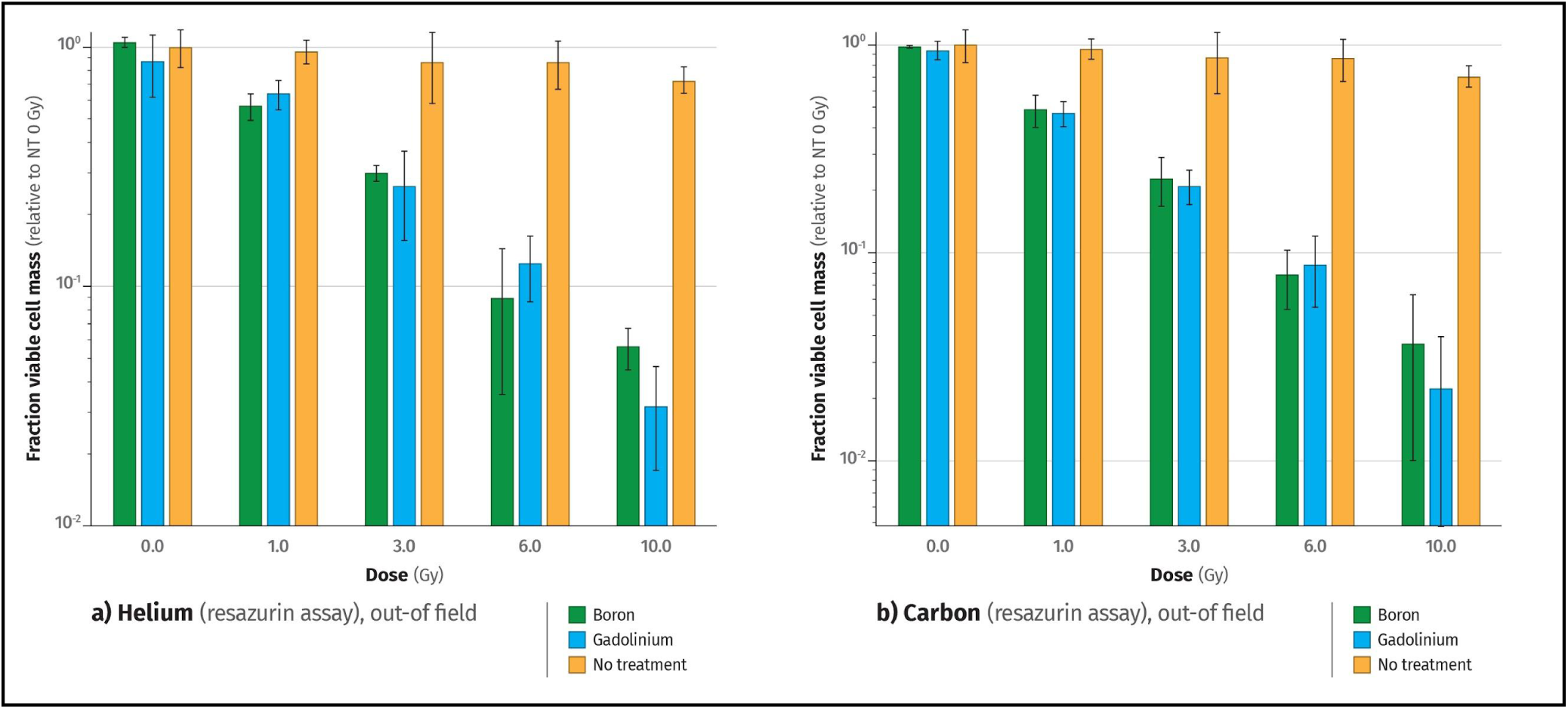
Out-of-field effect of neutron capture agents (NCAs) on cell survival. Fraction of viable cell mass (normalised to no-treatment, 0 Gy) for T98G cells positioned outside of the target volume, to the left and right of the centroid of the spread out Bragg peak region, irradiated with (a) helium or (b) carbon across a range of physical radiation doses, following incubation with 500 μM of either NCA, in triplicate. Even at the highest in-beam dose of 10 Gy, the no-NCA controls only see a decrease in relative viable cell mass of about 15%. By contrast, cells incubated with either NCA experience a decline in viability as the target dose is escalated, decreasing by more than 95% for a primary dose of 10 Gy. Bars represent the mean fraction viable cell mass of six independent experiments for the resazurin assay, and the T-bars represent ±2 standard deviations.

### Effect of NCA concentration

The effect of NCA concentration on cell viability with a fixed in-beam helium and carbon ion dose is illustrated in Figure 8. IC50 values for [^10^B]BPA with 3 Gy of carbon and helium ions were 9.33 μM and 30.04 uM, respectively. IC50 values for [^157^Gd]DOTA-TPP with 3 Gy of carbon and helium ions were 21.72 uM and 8.89 uM, respectively.

**Figure 8:**
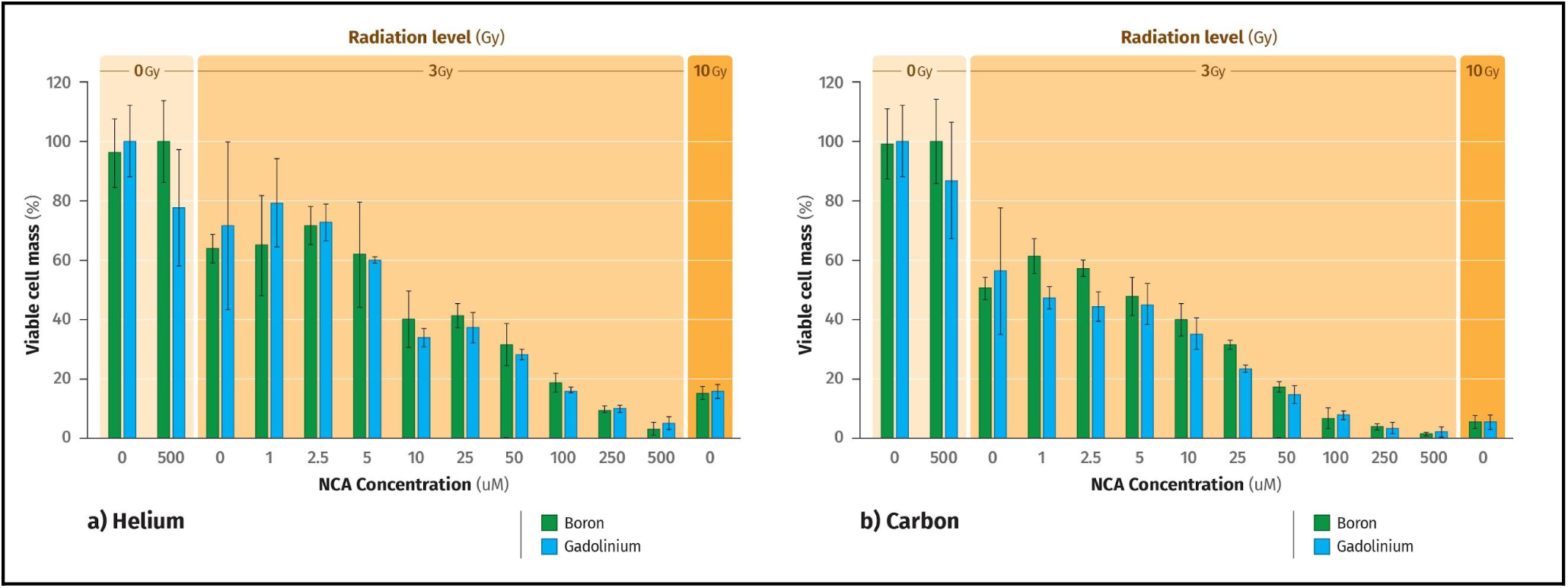
Concentration-dependent effect of neutron capture agents (NCAs) on cell viability. Viable cell mass (%) for T98G cells positioned in the centre of the spread out Bragg peak region irradiated with (a) helium or (b) carbon, receiving a physical dose of 3 Gy and incubated with increasing concentrations of each NCA, in triplicate. Non-irradiated controls with 0 or 500 μM of each NCA were included along with a 10 Gy positive control. For comparison, a dose of 3 Gy delivered to cells incubated with a concentration of 100-250 μM of either NCA is roughly equivalent to a 10 Gy dose with no NCA present (right-most group). Bars represent the mean percentage viable cell mass of two independent experiments for the resazurin assay, and the T-bars represent ±2 standard deviations.

In the absence of radiation, neither NCA is significantly cytotoxic in concentrations of up to 500 μM, although [^nat^Gd]DOTA-TPP shows mild cytotoxicity at higher concentrations (>500 μM) (Supplementary Material Section 5).

## Discussion

This paper presents the first experimental evidence quantifying the NCEPT effect in-vitro.

Thermal neutron fluence was quantified at a range of positions inside and around the target volume via gold activation, demonstrating the presence of a clinically relevant thermal neutron field - the key enabling factor for NCEPT. The shape of the neutron fluence distribution is similar to thatpreviously predicted via Monte Carlo simulations^10^, where the highest measured thermal neutron fluence is observed in the vicinity of the SOBP region before gradually falling off in all directions (Figure 4).

Human glioblastoma cells were placed both inside and immediately adjacent to a target volume inside a PMMA phantom, with and without the presence of two low-toxicity NCAs, [^10^B]BPA and [^157^Gd]DOTA-TPP^31,32^ (Figures 5-8). The target volume was irradiated with a 60 mm carbon or helium ion SOBP, both at a range of radiation doses with a fixed NCA dose, and secondly with a constant ion dose (corresponding to the ≅LD30 dose for cells not treated with NCAs) and a serial dilution of NCA. Cells placed inside the treatment volume reached 10% survival with the administration of 1.46 Gy of carbon ions or 1.67 Gy of helium ions with BPA, and 1.37 Gy of carbon ions and 1.44 Gy of helium ions with [^157^Gd]DOTA-TPP compared to 4.73 Gy of carbon ions and 6.53 Gy of helium ions with no NCA. This is a result of the combined impact of the dose delivered directly by the primary ion irradiation and the dose due to neutron capture by ^10^B or ^157^Gd. Therefore, dose enhancement to the treatment volume could be incorporated into treatment planning to either increase the effective dose to the tumour per ion delivered to the target or, where entrance dose is a concern, deliver an equivalent dose to the tumour with fewer ions. NCEPT will thus provide a wide range of potential options to the treating physician for either reducing off-target effects or increasing the dose which can be safely delivered to the target.

Cells placed *outside* and adjacent to the treatment volume only exhibit a decrease in growth in response to escalating dose to the target when treated with either of the evaluated NCAs. The magnitude of the reduction in viability of NCA-treated cells outside the SOBP is similar to that observed following irradiation of NCA-treated cells within the SOBP for both helium and carbon ions (Figure 5). Helium and carbon ions exhibit a lateral scattering of less than 2 mm at 11 cm depth in human tissue^41^. This superior dose conformity is demonstrated by the minimal response to escalating ion dose to target in cells that are not treated with an NCA, shown in Figure 7, while the neutron fluence at this location remains at 80-90% of the peak value (Figure 4) suggesting that the dominant factor in the reduction in cell viability is the neutron capture process.

Comparing the in-beam dose response, it is evident that cells treated with 500 uM concentration of NCA show a much stronger dose response compared to the no-NCA control group. Obtaining a detailed cell survival response at very low ion dose values (i.e., multiple steps in the 0-2 Gy dose range) presented experimental challenges. Modifying too many parameters at once, such as altering the attenuator to reduce dose rate, which also changes the beam quality, or extending the irradiation intervals for high dose regimes at low dose rates, would have compromised the experiment’s integrity. Furthermore, varying the beam current wasn’t a viable option due to limited beamtime at facilities like HIMAC.

To navigate these constraints, we varied the neutron capture dose by titrating the NCA while keeping the ion dose constant (Figure 8). Employing a dilution series of NCAs allowed us to elucidate the relative contribution of the neutron capture dose effectively, allowing us to evaluate cell viability in response to varying neutron doses at a finer resolution than would have been possible through coarse adjustments of the primary ion beam. For example, an NCA concentration of 10 uM used with a 3 Gy primary helium or carbon ion beam dose induces a 60% reduction of the viable cell mass, while a concentration of 100 uM with a 3 Gy primary beam dose results in a neutron capture effect equivalent to a 10 Gy dose of primary beam radiation (72% and 90% reduction in viable cell mass for helium and carbon ion beams, respectively).

While the linear-quadratic model provides an excellent fit for the no-NCA clonogenic cell survival results, it is clearly unable to provide a satisfactory fit for the NCA-treated cells. Since the α and β parameters of the linear-quadratic model are constrained to be non-negative, and the curve of best fit for the NCA-treated cells is concave up, the least-squares fit results in β = 0 in each case, such that the linear-quadratic model becomes a simple linear model. Due to the poor fit of this linear model across the full evaluated range of doses, to compute the 1% and 10% survival fraction doses we only consider the first four dose values, over which the logarithm of the survival fraction is approximately linear with respect to dose; while the fit remains poor, at least it allows an approximation of the 1% and 10% survival doses to be obtained. Relaxing the non-negativity constraint results in a better fit, but yields a nonsensical negative value for β. This implies that for NCEPT, the relation between survival fraction and dose is considerably more complex compared to photon or conventional charged particle therapy, and is incompletely described by the biophysical assumptions of the linear-quadratic model, a finding which supports previous criticism of the general validity of the LQ model in the literature ^42,43^.

In summary, our results indicate that NCEPT has the potential to greatly extend the utility of conventional charged particle therapies, not only by increasing the dose delivered to the primary target volume, but also by enabling the biochemically targeted delivery of radiation dose outside of conventionally delineated treatment areas. This capability could be particularly beneficial for treating micro-infiltrates and metastases that fall below the detection threshold of current molecular imaging techniques used in treatment planning.

Drawing on the principles to radionuclide therapy, we can envision a new class of radiopharmaceutical where the radiation emission within the biological system is only triggered by an external radiation source, at a time and location determined by the treatment plan. NCEPT is fundamentally different from traditional radiosensitisation techniques, as it introduces an entirely new radiation dose instead of merely amplifying a pre-existing one. Such an approach could deliver radiation more precisely to cellular targets while drastically reducing the dose to excretory pathways in comparison to current radionuclide therapeutics (which is a major limiting factor in such therapies). Future NCEPT agents should prioritise low inherent chemical toxicity and the targeted delivery of either ^10^B to specific cellular or tissue sites, or ^157^Gd to specific organelle or cellular targets.

We note that the observed dose-enhancement effect shown in our experiments is clearly not related to proton fusion for several reasons. Firstly, our radiation source is either ^12^C or ^4^He ions, and in both cases, no known projectile or target fragmentation reaction is able to generate protons in quantities sufficient to account for the observed results. Second, our target neutron capture isotopes, enriched ^10^B or ^157^Gd, are not conducive to beam-target fusion (and this would be the case even if a high-energy proton beam were used). Lastly, our findings - both in-beam and out-of-beam - are entirely consistent with the measured neutron fluence surrounding the target, corroborating the work of Manandhar et al. and Hosobuchi et al. who were unable to reproduce the purported proton-boron fusion dose enhancement results^16,17^, and the corresponding Monte Carlo simulation results of Jacobsen et al. and Khaledi et al.^13,14^, that predict negligible dose enhancement during proton irradiation of a ^11^B-loaded target. We note that our observed results further support the hypothesis that the previously observed proton-(natural)-boron dose enhancement effect is likely due to neutron capture by ^10^B, which constitutes approximately 19.8% of natural boron, since it is expected that similar (or even greater) thermal neutron fluences to those observed in these experiments would exist around the target when irradiated at depth.

The results obtained to date provide strong justification for proceeding to in vivo experimental exploration of NCEPT. The data suggests the potential for doses administered during particle therapies to be significantly reduced. Reductions in doses administered to patients can be expected to accompany comparable reductions in normal tissue complications and unwanted side-effects of radiation on sensitive organs. Additionally, this work provides further impetus to investigate the potential of NCEPT in proton therapy. Previous simulation studies indicated that neutron fluence during proton therapy should be even higher than for carbon ion therapy, which would imply that the impact of NCEPT may be even more significant there^10^. The experimental results presented in this work strongly support that hypothesis; we have shown that helium ion beams generate a substantially larger thermal neutron fluence for a given biological dose compared to carbon ions. Given the wider availability of proton therapy compared to heavy ion therapy, this would also expand the availability of NCEPT to a much wider range of potential patients and diseases.

## Conclusion

NCEPT represents a new paradigm in charged particle therapy. The combination of biochemically targeted neutron-capturing pharmaceuticals with conformally-targeted charged particle therapy - in particular, through exploitation of an internally generated secondary/tertiary radiation field - gives the possibility of attaining a remarkably high specificity of energy deposition within cells and tissues. The utility of neutron capture has been shown to extend beyond the margins of the conventional image targeted treatment volume and provides opportunity to target undiagnosed cancer micro-infiltrates that are otherwise left untreated. NCEPT highlights the potential of low molecular weight, high specificity small molecules, labelled with ^10^B, for a much broader range of radiotherapeutic and radiosensitisation applications - an area previously underexplored in comparison with heavier elements and large molecules or nanoparticles. We have demonstrated the potential to repurpose existing pharmaceuticals ([^10^B]BPA), as well as the potential for novel compounds ([^157^Gd]DOTA-TPP), to increase the efficacy of existing charged particle beams. The therapeutic exploitation of internally-generated neutrons by neutron capture agents is a process hitherto entirely unexplored by medicinal chemists, biologists and pharmacologists. We strongly feel that its potential for impacting clinical outcomes and improving quality of life following treatment warrants further investigation.

## Data Availability Statement

All data generated or analysed during this study are included in the paper and its Supplementary Information files, and available for download from the following public repository. https://bitbucket.org/mitra_safavi/ncept-in-vitro-data/

These include a complete database of the raw neutron fluence measurements and gold activity (per foil), and the Resazurin fluorescence measurements colony count.

## Code Availability Statement

All simulation and analysis code is available for download from the following public repository: https://bitbucket.org/mitra_safavi/ncept-in-vitro-data/

These include a full database of neutron fluence measurement, Monte Carlo simulation code & data analysis code, and the cell colony counting scripts used in the classification and identification of the cell colonies.

## Supporting information

Supplementary material including figures and tables.

## Acknowledgements

This work was supported by the Australian Nuclear Science and Technology Organisation (ANSTO) and the Japanese National Institutes for Quantum Science and Technology (QST). We would like to thank Gabriele Enge from the Wollongong Isotope Geochronology Laboratory (WIGL) for advice and technical support on ICP-MS analysis. We thank Hitomi Sudo and Atsushi Tsuji from the Institute for Quantum Medical Sciences (iQMS) for facilitating access to crucial analytical equipment for experiments conducted at QST. We thank Chris Dobie for reviewing the article prior to submission. We acknowledge NST Biosciences (ANSTO) for providing laboratory space and consumables for the development of the in vitro cell based models.Computer simulations were supported by the National Computing Merit Allocation Scheme (NCMAS) and the Multi-modal Australian Sciences Imaging and Visualisation Environment (MASSIVE). Invaluable support and advice was provided by the beam operators and technicians at the Heavy Ion Medical Accelerator (HIMAC) in Chiba, Japan . A.C. was beneficiary of an Australian Government Research Training Program Scholarship. The project received additional support from the Japan National Committee for the Union for International Cancer Control (UICC-Japan), the University of Sydney’s Drug Discovery Initiative (DDI) and the Australian Institute for Nuclear Science and Engineering (AINSE). Access to the Australian Centre for Neutron Scattering (ACNS) was enabled by the National Collaborative Research Infrastructure Strategy of Australia (NCRIS).

## Author Contributions

N.H., B.F., N.W, R.M., A.C., K.B., D.F., N.M, L.R., J.D., J.B., U.G., K.B., S.G., A.R. and M.S.N. contributed to the conception and/or design for this work. N.H., R.M., A.C., F.S., R.H., A.D., A.M. and M.S.N, contributed to collection of data. N.H., A.C. and M.S.N. contributed to analysis and interpretation of the data. N.H., M.S.N. and K.M. contributed to the drafting of the article. R.M., F.S., B.F., N.W., D.F., E.L., A.D. and A.R. were responsible for critical revision of the article. N.H., B.F., R.M., N.W., A.C., F.S., K.B., D.F., E.L., K.M., J.B., J.D., A.D., U.G., S.G., R.H., N.M., A.M., L.R., A.R. and M.S.N. approved the final version of the article to be published.

