## Supplementary material including figures and tables. for "Neutron capture enhances dose and reduces cancer cell viability in and out of beam during helium and carbon ion therapy"

**Supplementary Data Figure 1:** Experimental configuration at the HIMAC Biology beamline. Positions labelled as Z correspond to those described in Figure 3. The approximate beam path is overlaid in red and extends from the beam tube aperture on the right to the PMMA phantom on the left.

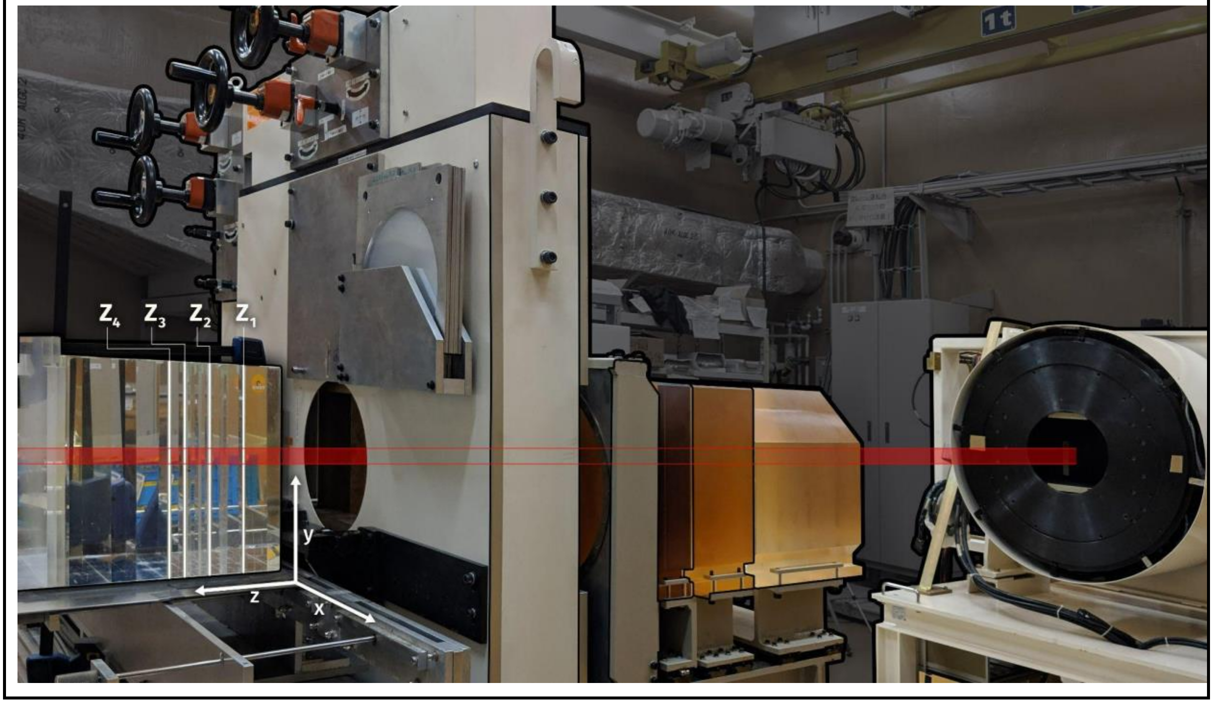


#### *Section 1*

##### Monte Carlo simulations of gold foil activation method for neutron fluence measurement

The accuracy of the proposed method for comparing differential activation of a pair of bare and cadmium-shielded $^{197}$Au samples for thermal neutron fluence estimation was established using Geant4 Monte Carlo simulations[1–3].

Geant4 10.2.p03 was used for all simulations in this study, with the selected physics models detailed in Supplementary Data Table 1. The choice of Geant4 version and physics models was based on a previously-published study of the accuracy of different Geant4 versions and hadronic inelastic ion fragmentation models by Chacon et al.[4].

**Supplementary Data Table 1:** Hadronic physics processes and models used in all simulations.

| **Interaction** | **Energy Range** | **Geant4 Model** |
|---|---|---|
| Radioactive Decay | All energies | G4RadioactiveDecayPhysics |

| | | |
|---|---|---|
| Particle Decay | All energies | G4Decay |
| Hadron Elastic | 0-100 TeV | G4HadronElasticPhysicsHP |
| Ion Inelastic | All energies | Binary Light Ion Cascade |
| Neutron Capture | < 20 MeV | NeutronHPCapture |
| | ≥ 20 MeV | nRadCapture |
| Neutron Inelastic | < 20 MeV | NeutronHPInelastic |
| | ≥ 20 MeV | Binary Cascade |

A 300×300×300 $mm^3$ PMMA phantom was irradiated with a polyenergetic circular (∅ = 100 mm) beam of $^{4}He$ or $^{12}C$ ions. Beam spectra were constructed so as to create a biologically flat spread-out Bragg peak extending between depths of 70 mm and 130 mm in the PMMA phantom, using the same procedure as described in a previous fragmentation model study by Chacon et al.[4]. A total of $2\times10^8$ primary particles were used for each simulation.

Bare/shielded detector pairs were placed at depths of 50 mm, 100 mm and 200 mm in the phantom, corresponding to the entrance, SOBP and tail regions. Three detectors pairs were positioned at each depth, with the first pair located at the centre of the beam, the second positioned at a 50 mm lateral offset from the centre (partially covered by the beam), and the third at a 120 mm lateral offset from the centre (outside of the beam, but still subject to some scattered primary particles from the beam). The shielded gold discs were placed at a 7 mm lateral lateral offset relative to the bare sample. The full set of locations is summarised in Supplementary Data Table 2.

**Supplementary Data Table 2:** Locations of gold samples within a 300 mm cubic PMMA phantom centred at coordinates (0, 0, 0).

| **Sample** | **Position** | **X (cm)** | **Y (cm)** | **Z (cm)** |
|---|---|---|---|---|
| Bare | 1 | 0 | -0.5 | 5 |
| | 2 | 5.5 | -0.5 | 5 |
| | 3 | 12.5 | -0.5 | 5 |
| | 4 | 0 | 0 | 10 |
| | 5 | 5.5 | 0 | 10 |

| | | | | |
|---|---|---|---|---|
| | 6 | 12.5 | 0 | 10 |
| | 7 | 0 | 0.5 | 15 |
| | 8 | 5.5 | 0.5 | 15 |
| | 9 | 12.5 | 0.5 | 15 |
| Shielded | 1 | 0 | -1.2 | 5 |
| | 2 | 5.5 | -1.2 | 5 |
| | 3 | 12.5 | -1.2 | 5 |
| | 4 | 0.7 | 0 | 10 |
| | 5 | 6.5 | 0 | 10 |
| | 6 | 6.5 | 0 | 10 |
| | 7 | 13.2 | 1.2 | 15 |
| | 8 | 5.5 | 1.2 | 15 |
| | 9 | 12.5 | 1.2 | 15 |

Photons with energies between 410 keV and 413 keV emitted from the gold between 6 h and 144 h post-irradiation were scored, along with the parent radionuclide which created them (the 6 h minimum being required to allow for activation products other than $^{198}$Au to decay). The fraction of gamma photons originating from the decay of $^{198}$Au was estimated; in addition, the process which resulted in the creation of all $^{198}$Au nuclei and the total number of thermal neutrons which entered into the gold samples (used to estimate the ground truth neutron fluence) at each location was also recorded. This combination allows the effects of the ion beam on the accuracy of neutron activation analysis to be evaluated, since any activations resulting in the emission of gamma photons in the 410 keV-413 keV range that are not a result of thermal neutron capture can be quantified, and the estimate based on relative activation of the shielded and unshielded gold detectors can be compared directly with the ground truth thermal neutron fluence.

Supplementary Data Tables 3 and 4 show, for helium and carbon irradiation, respectively, the mean number of activations in both the bare and shielded gold foils, the estimated thermal neutron fluence $\mathbf{\Phi_{th}}$. The percentage error was calculated against ground truth fluence.

| **Supplementary Data Table 3:** Neutron fluence estimates during simulated helium ion irradiation (after correcting for field isotropy), together with ground truth and relative percentage error. | | | | | |
|---|---|---|---|---|---|
| **Position** | **$^{198}$Au (bare)** | **$^{198}$Au (shielded)** | **$\Phi_{th}$ (estimated)** | **$\Phi_{th}$ (ground truth)** | **Error (%)** |
| 1 | 668 | 153 | 83,300 | 86,700 | +3.95 |
| 2 | 477 | 100 | 61,000 | 63,700 | +4.25 |
| 3 | 123 | 23.2 | 16,200 | 17,800 | +9.16 |
| 4 | 825 | 165 | 107,000 | 106,000 | -0.68 |
| 5 | 608 | 97.4 | 82,600 | 80,100 | -3.16 |
| 6 | 165 | 26.4 | 22,500 | 22,900 | +1.85 |
| 7 | 393 | 56.8 | 54,400 | 54,100 | .0.49 |
| 8 | 432 | 38.0 | 63,700 | 60,900 | -4.61 |
| 9 | 151 | 10.2 | 22,800 | 20,100 | -13.3 |

| **Supplementary Data Table 4:** Neutron fluence estimates during simulated carbon ion irradiation (after correcting for field isotropy), together with ground truth and relative percentage error. | | | | | |
|---|---|---|---|---|---|
| **Position** | **$^{198}$Au (bare)** | **$^{198}$Au (shielded)** | **$\Phi_{th}$ (estimated)** | **$\Phi_{th}$ (ground truth)** | **Error (%)** |
| 1 | 150 | 31 | 19,300 | 19,500 | 0.99 |
| 2 | 117 | 25 | 14,900 | 14,800 | -0.73 |
| 3 | 30.6 | 4.2 | 427 | 4,440 | 3.86 |
| 4 | 202 | 35.6 | 26,900 | 26,100 | -2.97 |
| 5 | 154 | 27.4 | 20,500 | 20,000 | -2.52 |
| 6 | 41.2 | 8.8 | 5,240 | 5,880 | 10.9 |
| 7 | 131 | 21.6 | 17,700 | 184,000 | 3.86 |
| 8 | 131 | 19.6 | 18,000 | 17,600 | -2.16 |
| 9 | 41 | 5.4 | 5,760 | 6,000 | 4.06 |

In all locations, more than 99.9% of gamma photons with energies between 410 keV and 413 keV were a result of neutron capture in the gold foil; in addition, almost zero gamma photons in this range originated from the cadmium shield. This, combined with the excellent agreement of the estimated thermal neutron fluence based on the differential activation of the bare and shielded gold foils with the ground truth strongly confirm the theoretical basis for the use of the gold foil activation method for quantifying thermal neutron fluence in the high-intensity mixed radiation field present during helium or carbon ion therapy.

#### Experimental quantification of thermal neutron fluence by gold foil activation

The thermal neutron fluence at each point of measurement resulting from irradiation of the PMMA target with helium ions are listed in Table 5; corresponding results for carbon are shown in Table 6. The mean thermal neutron fluence at the centre of the SOBP is approximately $5.8\times10^9$ n/cm2/Gy(RBE) for the helium beam and $2.2\times10^9$ n/cm2/Gy(RBE) for the carbon beam, respectively.

**Supplementary Data Table 5:** Thermal neutron yields measured in the PMMA phantom using helium SOBP

| **Depth** | **Plate Rotation (Degrees)** | **Foil Letter** | **Thermal Neutron fluence n/(cm$^2$ Gy(RBE))** |
|---|---|---|---|
| Entrance 48mm | 0 | A | $(5.7 \pm 0.2)$ x$10^9$ |
| | | B | $(5.1 \pm 0.2)$ x$10^9$ |
| | | C | $(3.4 \pm 0.2)$ x$10^9$ |
| | | D | $(4.8 \pm 0.2)$ x$10^9$ |
| | | E | $(2.1 \pm 0.1)$ x$10^9$ |
| Middle SOBP 100mm | 90 | A | $(6.2 \pm 0.3)$ x$10^9$ |
| | | B | $(5.7 \pm 0.3)$ x$10^9$ |
| | | C | $(3.7 \pm 0.2)$ x$10^9$ |
| | | D | $(5.2 \pm 0.3)$ x$10^9$ |
| | | E | $(2.4 \pm 0.2)$ x$10^9$ |
| | | A | $(5.4 \pm 0.3)$ x$10^9$ |

| | | | |
|---|---|---|---|
| Distal SOBP 133mm | 180 | B | (4.8 ± 0.3) x$10^9$ |
| | | C | (3.4 ± 0.2) x$10^9$ |
| | | D | (4.6 ± 0.3) x$10^9$ |
| | | E | (2.4 ± 0.4) x$10^9$ |
| Tail 153mm | 270 | A | (5.1 ± 0.3) x$10^9$ |
| | | B | (4.7 ± 0.3) x$10^9$ |
| | | C | (3.0 ± 0.2) x$10^9$ |
| | | D | (4.15 ± 0.2) x$10^9$ |
| | | E | (2.0 ± 0.5) x$10^9$ |

**Supplementary Data Table 6:** Thermal neutron yields measured in the PMMA phantom using carbon SOBP

| **Depth** | **Plate Rotation (Degrees)** | **Foil Letter** | **Thermal Neutron fluence n/(cm$^2$ Gy(RBE))** |
|---|---|---|---|
| Entrance 48mm | 0 | A | (2.1 ± 0.1) x$10^9$ |
| | | B | (1.9 ± 0.1) x$10^9$ |
| | | C | (1.4 ± 0.1) x$10^9$ |
| | | D | (1.7 ± 0.1) x$10^9$ |
| | | E | (0.8 ± 0.2) x$10^9$ |
| Middle SOBP 100mm | 90 | A | (2.3 ± 0.2) x$10^9$ |
| | | B | (1.9 ± 0.2) x$10^9$ |
| | | C | (1.3 ± 0.2) x$10^9$ |

| | | | |
|---|---|---|---|
| | | D | (1.8 ± 0.1) x$10^9$ |
| | | E | (0.98 ± 0.09) x$10^9$ |
| Distal SOBP 133mm | 180 | A | (2.2 ± 0.2) x$10^9$ |
| | | B | (2.2 ± 0.1) x$10^9$ |
| | | C | (1.6 ± 0.2) x$10^9$ |
| | | D | (2.0 ± 0.1) x$10^9$ |
| | | E | (0.80 ± 0.09) x$10^9$ |
| Tail 153mm | 270 | A | (2.0 ± 0.3) x$10^9$ |
| | | B | (2.0 ± 0.2) x$10^9$ |
| | | C | (1.2 ± 0.1) x$10^9$ |
| | | D | (1.4 ± 0.1) x$10^9$ |
| | | E | (1.0 ± 0.1) x$10^9$ |

The shape of the neutron fluence distribution is similar to those predicted in the carbon ion Monte Carlo simulations, where the highest measured thermal neutron fluence is in the vicinity of the SOBP region before gradually falling off in all directions. However, the magnitude of the measured field is considerably higher. There are several possible reasons for this discrepancy, including the use of a passively scattered SOBP rather than an active beam delivery system, possible beamline contamination (which would principally manifest as fast neutrons, some of which would thermalise in the target), and the choice of simulation physics models (which may not properly reflect the true rate of neutron production). Further simulation and experimental study of the thermal neutron component of the radiation field in heavy ion therapy is therefore warranted. Regardless of the reasons for the larger than expected magnitude of the thermal neutron field, its presence is beneficial for NCEPT as it will proportionally increase the dose which may be delivered via thermal neutron capture.

### *Section 2.*

#### Derivation of self shielding ratio

The self-shielding term $G_{th}$ is calculated according to the method described by Fleming[5]. Neglecting internal scatter, the self-shielding factor for a flat slab of thickness $T$ is given by

$$G_{th-NS} = \frac{1}{x}\left[\frac{1}{2} - E_3(x)\right]$$

where $x = T\Sigma_a$ for a slab of thickness $T$, and $E_3(x) = \int_1^{\infty} \frac{e^{-xt}}{t^3} dt$

If the neutron spectrum follows a Maxwellian energy distribution (as it is assumed to do in a thermalised neutron field), $\Sigma_a$ is modified[6]:

$$\Sigma_a = \frac{2}{\sqrt{\pi}}\sqrt{\frac{T_0}{T_n}}\Sigma_a(v_0)$$

where $T_0$ = 293.6 K and $v_0$ = 2200 ms$^{-1}$. Finally, to account for internal neutron scattering, the shielding factor must be adjusted[5]:

$$G_{th} = \frac{G_{th-NS}}{1 - \frac{\Sigma_s}{\Sigma_t}(1 - G_{th-ns})}$$

where $\Sigma_s$ is the macroscopic scattering cross-section and $\Sigma_t$ is the macroscopic total cross-section (0.428 cm$^{-1}$ and 6.26 cm$^{-1}$, respectively, for $^{197}$Au. For the dimensions of gold foils used in the simulation, the isotropic-field Maxwellian-spectrum self-shielding factor is 0.96494.

### *Section 3.*

#### Synthesis, purification and preparation of NCA's

##### Preparation of [$^{10}$B]BPA

[$^{10}$B]BPA (>98.4% $^{10}$B) was purchased from Interpharma Praha, a.s., (Prague, Czech Republic). [$^{10}$B]BPA was dissolved directly into Eagles Minimum Essential Media (EMEM) at a concentration of 500 µM by incubating at 40 °C while agitating in an ultrasonic water bath for 30-40 min.

##### Synthesis of [$^{157}$Gd]DOTA-TPP

[$^{157}$Gd]DOTA-TPP was synthesised following a modified two-step synthesis based upon the work of Morrison *et al*[7]. In the first reaction step, the reagents (4-(bromomethyl)benzyl)triphenyl phosphonium bromide (460 mg, 0.87 mmol), tri-*tert*-butyl

1,4,7,10-tetraazacyclododecane -1,4,7-triacetate (530 mg, 1.03 mmol) and $NaHCO_3$ (140 mg, 1.67 mmol) were added to a 25 mL round bottom flask and dissolved up in 4 mL of anhydrous acetonitrile. The reaction mixture was then stirred and heated to reflux (82 °C) for 4 h and then cooled to room temperature and filtered through a 0.45 µm syringe filter (~4 mL acetonitrile + 2×2 mL acetonitrile wash). The acetonitrile from the filtrate was then removed by evaporation under reduced pressure and the resulting solid was dissolved up in 5 mL of dichloromethane. To this solution, 5 mL of trifluoroacetic acid was added; it was then stirred at room temperature for a further 20 h. The excess trifluoroacetic acid (~5 mL) was then removed by passage of a stream of nitrogen gas over the solution for 12 h. The resulting brown solid was then purified by preparative reverse phase HPLC using an Atlantis T3 OBD C18 150×30 mm, 10 µm column, eluting with 20% $C_2H_3N$/75% $H_2O$/5% $NH_4OAc$ (0.1 M, pH 4.3) at 30 mL/min ($t_R$ = 8.5-12 min); $\lambda$=254 nm (see extended data). The fractions containing the ligand were combined and the excess acetonitrile removed by evaporation under reduced pressure. The remaining water and excess $NH_4OAc$ was then removed by lyophilisation to give the free ligand DOTA-TPP.OAc as a white solid (374 mg, 40%). $^1H$ NMR ($D_2O$) δ 7.82-7.78 (m, 3H), 7.63-7.53 (m, 12H), 7.29 (d, *J* = 7.8 Hz, 2H), 6.94 (q, *J* = 1.7, 7.8 Hz, 2H), 4.78 (s, 10H), 4.72 (s, 1H), 4.69 (s, 1H), (3.76, bs, 2H), 3.53-2.79 (m, 22H), 1.90 (s, 5H). $^{13}C$ NMR ($D_2O$) δ 180.76, 176.61, 171.38, 135.81, 135.79, 134.65, 134.56, 132.42, 132.37, 131.69, 131.67, 130.66, 130.54, 128.05, 127.98, 57.31, 56.71, 56.12, 51.59, 50.92, 49.21, 48.45, 30.15, 29.67, 23.27. LRMS: predicted for $C_{40}H_{48}N_4O_6P^+$ = 711.33, found = 711.5 ($M^+$), 356.4 (($M^+$ + $H^+$)/2). HPLC purity = 99.3% using an Atlantis T3 150×4.8 mm, 5 µm. eluting with a gradient of 5% $C_2H_3N$/85% $H_2O$/10% aq.TFA (1%) → 90% $C_2H_3N$/10% aq.TFA (1%) over 20 min at 1 mL/min ($t_R$ = 11.5 min); $\lambda$=254 nm (see extended data). Elemental analysis: Predicted for (DOTA-TPP.OAc).1.2(AcOH).$NH_4OAc$ = C 60.58, H 6.88, N 7.61. Found: C 59.88, 60.42, H 6.61, 7.03, N 7.43, 7.47. For the second reaction step the DOTA-TPP free ligand (770 mg, 0.84 mmol) was dissolved in 20 mL of water and [$^{157}$Gd]$Gd_2O_3$ (218 mg, 0.60 mmol) was added. The suspension was stirred and heated to 80 °C for 20 h and then the unreacted [$^{157}$Gd]$Gd_2O_3$ was removed by passage through a 0.45 µm syringe filter (~20 mL water + 2×2 mL water wash). The water was then removed by evaporation under reduced pressure to give the final compound [$^{157}$Gd]DOTA-TPP as a colourless solid (881 mg, quantitative yield). HPLC purity = 96% using an Atlantis T3 150×4.8 mm, 5 µm. eluting with a gradient of 10% $C_2H_3N$/80% $H_2O$/10% aq.TFA (1%) → 90% $C_2H_3N$/10% aq.TFA (1%) over 20 min at 1 mL/min ($t_R$ = 10.5 min); $\lambda$=254 nm. LRMS: predicted for enriched $C_{40}H_{45}{}^{157}GdN_4O_6P^+$ (90.4% Gd-157, 6.3% Gd-158, 2.65% Gd-156, 0.5% Gd-160, 0.1% Gd-155, 0.05% Gd-154) = 865.35, found = 865.52 ($M^+$). Elemental analysis: Predicted for [$^{157}$Gd]DOTA-TPP.OAc.7$H_2O$ = C 47.99, H 5.94, N 5.33. Found: C 47.75, 47.97, H 5.58, 5.54, N 5.11, 5.03.

##### Preparative HPLC conditions for DOTA-TPP

Preparative reverse phase HPLC conditions: The DOTA-TPP free ligand was purified on a WATERS preparative HPLC system consisting of a W25X5Q quaternary gradient solvent pump, W2707 autosampler, W2489 UV/Visible detector set at $\lambda$=254 nm and a WFC-III fraction collector. The compound was purified on an Atlantis T3 OBD C18 150×30 mm, 10 µm column, eluting with 20% $C_2H_3N$/75% $H_2O$/5% $NH_4OAc$ (0.1 M, pH 4.3) at 30 mL/min ($t_R$ = 8.5-12 min).

##### QC HPLC conditions for DOTA-TPP

Quality Control (QC) reverse phase HPLC conditions: The final DOTA-TPP free ligand was analysed on a WATERS QC HPLC system consisting of a W600 quaternary gradient solvent pump, W2717 autosampler, W2996 PDA UV detector set at $\lambda$=254 nm and injected on an

Atlantis T3 150×4.8 mm, 5 µm column. eluting with a gradient of 5% $C_2H_3N$/85% $H_2O$/10% aq.TFA (1%) → 90% $C_2H_3N$/10% aq.TFA (1%) over 20 min at 1 mL/min ($t_R$ = 11.5 min); $\lambda$=254 nm.

### Section 4.

#### Radiobiological assays

##### Growth Curve Data from Cell Growth Assay

<table>
<tr><td colspan="4">Supplementary Data Figure 2: Representative growth curves obtained from monitoring cell growth via resazurin assay following irradiation with a dose escalation of carbon or helium ions. Cells were treated with either [10]B-BPA or [157Gd]-Gd-DOTA-TPP at 500uM.</td></tr>
<tr><td></td><td>Carbon</td><td>Helium</td><td rowspan="4">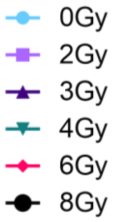</td></tr>
<tr><td>[10B]-BPA 500uM</td><td>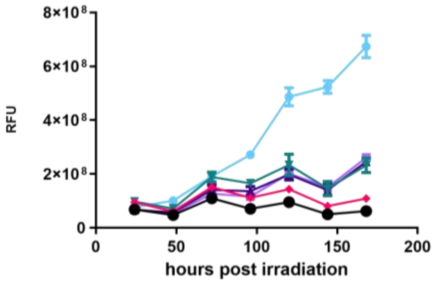</td><td>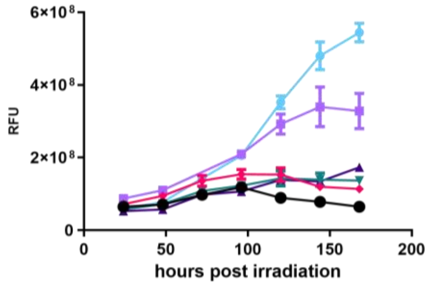</td></tr>
<tr><td>[157Gd]-GdTPP 500uM</td><td>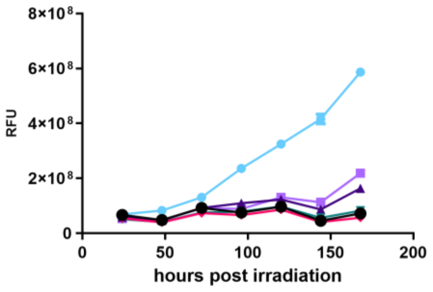</td><td>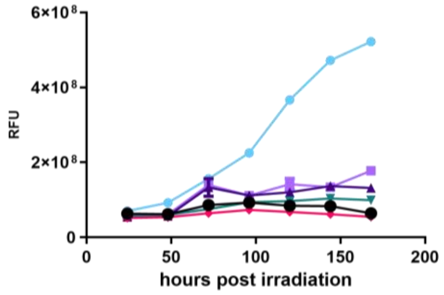</td></tr>
<tr><td>No Treatment</td><td>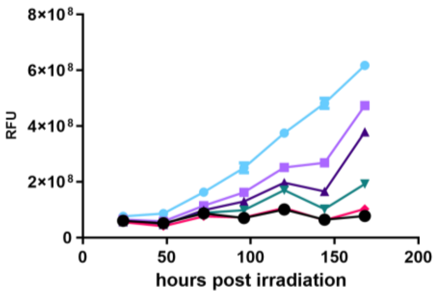</td><td>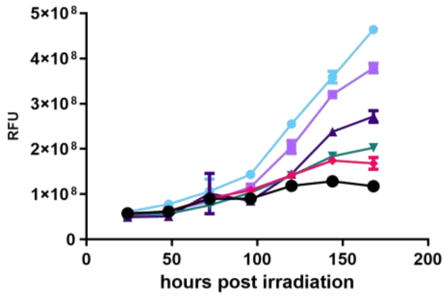</td></tr>
</table>

**Supplementary Data Figure 3:** Representative growth curves obtained from monitoring cell growth via resazurin assay following irradiation of the primary target volume with a dose escalation of carbon or helium ions and the cells placed outside and immediately adjacent to the target. Cells were treated with either [10]B-BPA or [157Gd]-Gd-DOTA-TPP at 500uM.

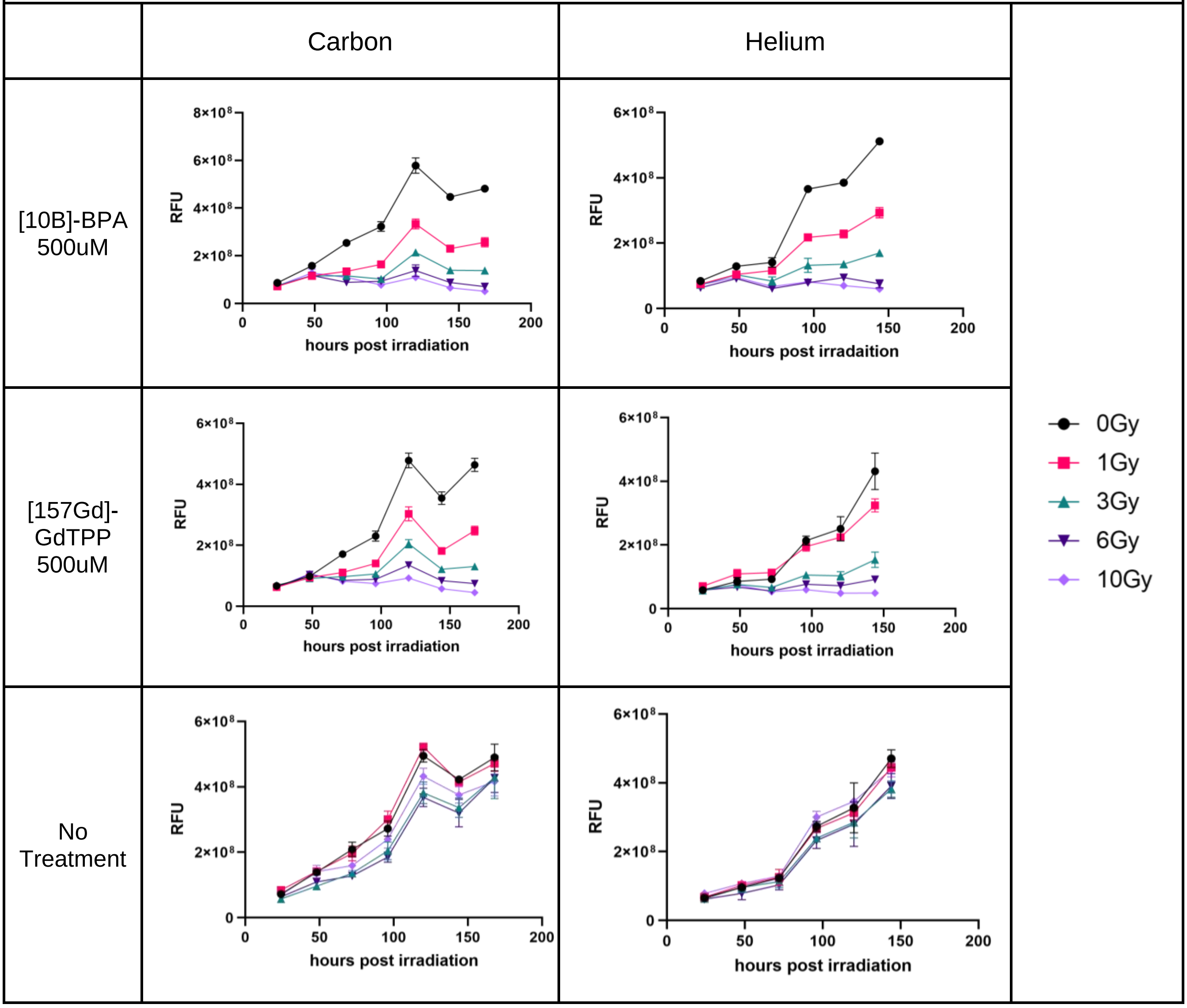


#### Seeding Densities for Clonogenic Assay

**Supplementary Data Table 7:** Seeding density (cells / Petri dish) of the Petri dishes for the clonogenic assay.

| **Dose (Gy)** | **Control** | **[$^{10}$B]-BPA** | **[$^{157}$Gd]DOTA-TPP** |
|---|---|---|---|
| 0 | 3,000 | 5,000 | 7,000 |
| 2 | 6,000 | 9,000 | 12,000 |
| 3 | 7,000 | 10,000 | 13,000 |

| | | | |
|---|---|---|---|
| 4 | 15,000 | 60,000 | 100,000 |
| 6 | 30,000 | 60,000 | 120,000 |
| 8 | 40,000 | 80,000 | 120,000 |
| 10 | 40,000 | 100,000 | 150,000 |

#### Clonogenic Assay LQ model

All colony count data and images stored at [https://bitbucket.org/mitra_safavi/ncept-in-vitro-data/src/master/Clonogenic/](https://bitbucket.org/mitra_safavi/ncept-in-vitro-data/src/master/Clonogenic/)

**Supplementary Data Table 8:** α and β parameters of the linear quadratic model fitted to the survival fraction obtained via the clonogenic assay together with the α : β ratio.

| **Compound** | **Carbon** | | | **Helium** | | |
|---|---|---|---|---|---|---|
| | *α* ($Gy^{-1}$) | *β* ($Gy^{-2}$) | *α:β (Gy)* | *α* ($Gy^{-1}$) | *β* ($Gy^{-2}$) | *α:β (Gy)* |
| Control | 0.343 | 0.0304 | 11.3 | 0.132 | 0.0338 | 3.91 |
| [$^{10}$B]-BPA | 1.58 | 0 | ∞ | 1.37 | 0 | ∞ |
| [$^{157}$Gd]-DOTA-TPP | 1.68 | 0 | ∞ | 1.59 | 0 | ∞ |

#### Assessment of a selection of automated colony counting methods

Three open source image based colony counting methods were assessed for the ability to determine colony forming units (CFU) by comparing the software outputs from digitised petri dishes with the mean output of three human operators on the same randomised selection of clonogenic assay samples. The software packages assessed were:

- Open CFU ([https://github.com/qgeissmann/OpenCFU](https://github.com/qgeissmann/OpenCFU))
- Analyse Particles - ImageJ ([https://imagej.net/imaging/particle-analysis#analyze-particles](https://imagej.net/imaging/particle-analysis#analyze-particles))
- Colony Counter ([https://imagej.nih.gov/ij/plugins/colony-counter.html](https://imagej.nih.gov/ij/plugins/colony-counter.html))

The outputs of each program were plotted against the mean counts of the human operators and a straight line fitted to the data. The resulting $R^2$ values indicate the degree of agreement between the software method and the human operator.

**Supplementary Data Figure 4**:
Plots of manual(operator) counts versus software outputs for colony forming units on petri dishes

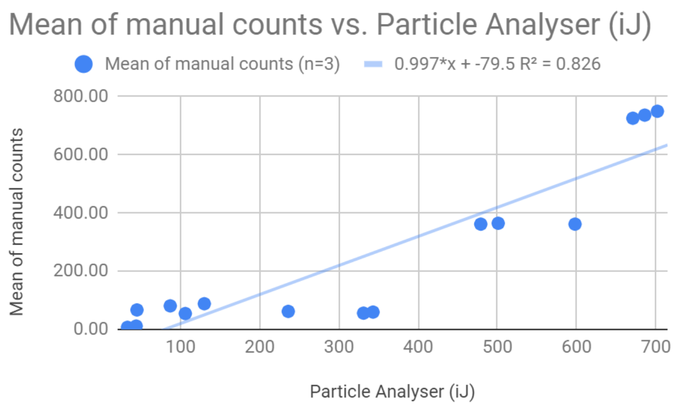


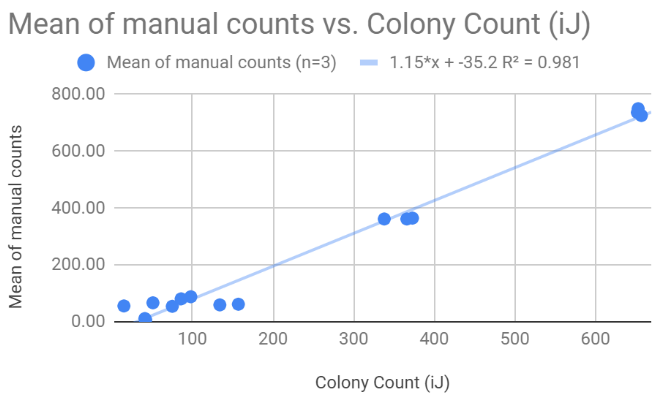


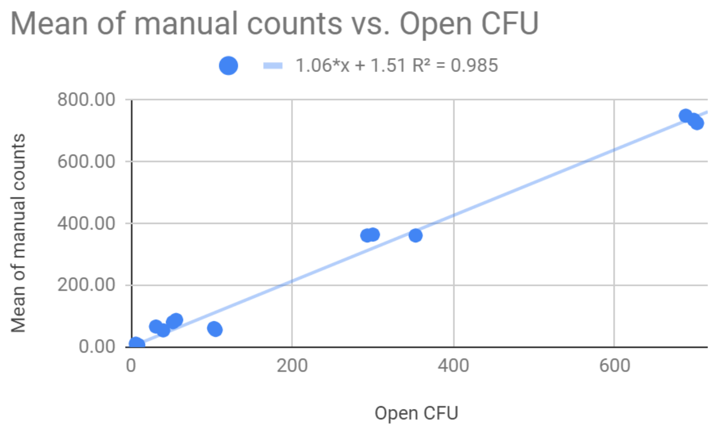


For these samples, Open CFU showed the highest level of agreement to the human operators with an $R^2$ of 0.985, followed by Colony Counter and Particle Analyser. This criteria was used to select Open CFU as the method used for batch processing data from all petri dishes produced for the clonogenic assay.

### Section 5.

#### Compound cytotoxicity and uptake

The cytotoxic and cytostatic activity of BPA and [natGd]DOTA-TPP on T98G cultures was measured by resazurin cell viability assay[8]. T98G cells were seeded into a 96 well plate at a density of 5,000 cells per well and allowed to attach overnight. Cells were treated, in triplicate, with a serial dilution (1,000, 500, 100, 50, 10, 5, 1, 0.1 µM) of each NCA and a serial dilution (500, 100, 10, 1, 0.1, 0.01, 0.001, 0.0001 µg/mL) of Digitonin (Sigma-Aldrich) in complete EMEM. Cells were placed into the incubator for 24 h at which time the relative viable cell mass in each well was measured by resazurin cell viability assay as follows: A 1%w/v stock solution of resazurin sodium salt was prepared in water. The stock was diluted 250 times in phosphate-buffered saline (PBS) and warmed to 37 °C in a water bath. All media was removed from the plate, replaced with resazurin-PBS and incubated for 60 min at 37 °C. The resulting conversion of resazurin to resorufin was measured by recording the fluorescence at excitation wavelength 555 nm and emission wavelength 585 nm at 37 °C. The resulting background subtracted relative fluorescence units (RFU) were plotted against log-concentration and an IC50 value determined by fitting a sigmoidal log-inhibitor vs response model (Supplementary Data Figure 2).

All data available at https://bitbucket.org/mitra_safavi/ncept-in-vitro-data/src/master/Cytotox/

**Supplementary Data Figure 5:**

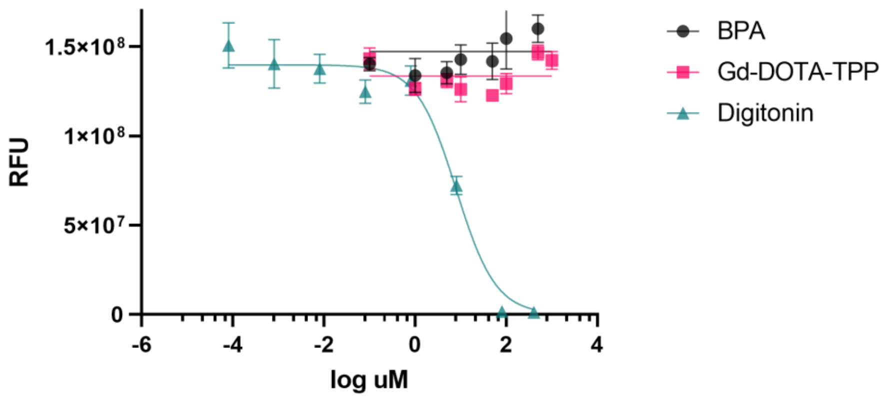


| | BPA | Gd-DOTA-TPP | Digitonin |
|---|---|---|---|
| IC50 (µM) ± SEM | NC | NC | 9.554 ± 1.556 |

*NC = not calculated

Cytostatic and cytotoxic potential of BPA and Gd-DOTA-TPP on T98G tissue cultures compared to Digitonin. No measurable effect was observed for either neutron capture agent and no IC50 was able to be calculated compared to 9.554 µM for Digitonin.

T98G cells were incubated, in triplicate, for 24 h in complete growth media, or complete growth media containing either 500 µM BPA or 500 µM [$^{nat}$Gd]DOTA-TPP. The cells were washed thoroughly in PBS, lysed in 400ul of 65% nitric acid (Suprapur, Merk) and then left to digest at room temperature for 48 h, with gentle agitation. Total protein content of the digests was determined by assaying the increase in absorbance of each sample at 358 nm relative to similarly digested standards of varying concentrations of bovine serum albumin. The resulting cell digests were transferred to 50ml falcon tubes and diluted 1/50 with MilliQ water. B and Gd elemental concentration analyses conducted on a quadrupole inductively-coupled plasma mass spectrometer (ThermoFisher iCAP-Q™) at the Wollongong Isotope Geochronology Laboratory (WIGL), University of Wollongong (Supplementary Data Table 9). The sample introduction system consisted of a PFA nebuliser, quartz spray chamber, nickel sample and skimmer cones fitted with a high-matrix insert. At the start of the session, the instrument was tuned with ThermoFisher tuning solution. Calibration standards and samples were introduced into the instrument as a 0.3M $HNO_3$ solution made from ultra-trace grade $HNO_3$ (Seastar Baseline™ grade). Analytes $^{11}$B and $^{157}$Gd were collected with a dwell time of 0.1 s for each over 90 sweeps. Quantification was performed by analysing a set of calibration standards at the start of the analysis sequence, with concentrations ranging from 0.0001 to 25 ppb. The calibration curve was determined using a linear regression through the calibration standards and forcing through the blank. The correlation coefficient for the calibration curve was 1.0 for both analytes. The limit of detection was 0.008 ppb for B and 0.0001 ppb for Gd. Relative standard error on the calibration curve was 14% for B and 5.7% for Gd. Blanks were measured during analysis and were <0.05 ppb for B and <0.001 ppb for Gd. Accuracy during analysis was tested by measuring 1 ppb calibration standard as an unknown, and it was determined to be 8% for B and 1.3% for Gd. The mass of Gd or B in the digests, along with the protein concentration, were used to calculate the mass of NCA per mg of protein, and then finally average mass of NCA per cell (Supplementary Data Table 10).

**Supplementary Data Table 9**: ICP-MS data, given in parts per billion (ppb), alongside the mass of total protein (µg) determined for each individual digest by nitric acid protein assay.

| | **Total B or Gd (ppb)** | **Protein (µg)** |
|---|---|---|
| BPA | 5.57 ± 0.28 | 116.64 |
| | 3.64 ± 0.08 | 123.58 |
| | 5.63 ± 0.5 | 135.83 |
| BPA negative | 0.64 ± 0.01 | 116.24 |
| | 0.20 ± 0.01 | 128.63 |

| **Supplementary Data Table 9**: ICP-MS data, given in parts per billion (ppb), alongside the mass of total protein (µg) determined for each individual digest by nitric acid protein assay. | | |
|---|---|---|
| | 0.35 ± 0.01 | 130.84 |
| B Blank | <0.1 ± 0.01 | - |
| Gd-DOTA-TPP | 10.29 ± 0.02 | 142.58 |
| | 9.71 ± 0.1 | 148.87 |
| | 9.98 ± 0.03 | 102.96 |
| Gd-DOTA-TPP negative | <0.01 ± 0.01 | 83.27 |
| | <0.01 ± 0.01 | 91.72 |
| | <0.01 ± 0.01 | 301.61 |
| Gd Blank | <0.01 ± 0.01 | - |

| **Supplementary Data Table 10**. B and Gd quantities per cell | | |
|---|---|---|
| | **atoms/cell** | **SD** |
| **Boron** | $4.72 \times 10^{8}$ | $1.12 \times 10^{8}$ |
| **Gadolinium** | $6.97 \times 10^{7}$ | $1.49 \times 10^{7}$ |

### Section 6.

#### Neutron capture effect of non-enriched Gd-DOTA-TPP

| **Supplementary Data Figure 6:** |
|---|

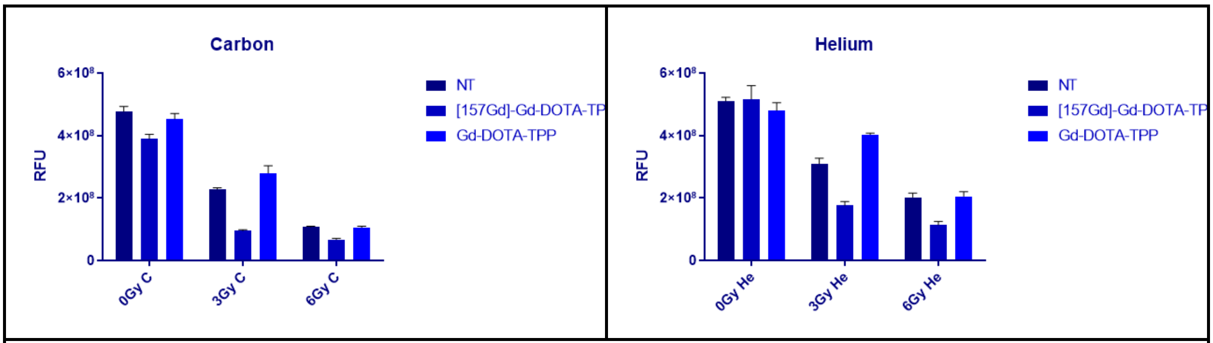


Effect of naturally abundant Gd labelled Gd-DOTA-TPP on T98G cell viability at day 6 following irradiation with Carbon or Helium beams. Compared with response for [$^{157}$Gd]-DOTA-TPP and no treatment (NT) control under the same conditions.
